# AT-HOOK-MOTIF NUCLEAR LOCALIZED 15 extends plant longevity by binding at poorly accessible, epigenetic mark-depleted chromatin surrounding transcribed regions

**DOI:** 10.1101/2025.08.05.668696

**Authors:** Thalia Luden, Jihed Chouaref, Remko Offringa

## Abstract

**Background:** Members of the *AT-HOOK MOTIF NUCLEAR LOCALIZED (AHL)* gene family have been shown to play important roles in plant development. In *Arabidopsis thaliana*, one member of this family, *AHL15*, induces somatic embryogenesis and extends plant longevity when overexpressed - the latter through strong repression of several ageing-related developmental transitions. However, its direct target genes and the mechanisms by which it regulates their expression have remained elusive to date.

**Results:** In this study we identified the genome-wide DNA binding sites of AHL15 and show that AHL15 binds throughout the genome at AT-rich sequences near the transcription start- and end sites in regions depleted of epigenetic marks. We show that induction of AHL15 activity causes strong and rapid changes in transcription, with the majority of the differentially expressed genes being downregulated but without directly affecting chromatin accessibility, resulting in developmental defects. In addition, AHL15 binding to regions near the transcription start and end sites was enhanced at genes that were differentially expressed upon AHL15 induction and was especially strong near the transcription start site of upregulated genes and near the transcription end site of downregulated genes. Finally, we show that AHL15 shares binding sites with the chromatin architectural protein GH1-HMGA2/HON5, which was previously shown to alter transcription by disrupting gene loop formation.

**Conclusions:** Together, our findings suggest that AHL15 affects the expression of its target genes by regulating the 3D organization rather than by changing the accessibility of chromatin or the deposition of histone modifications.

## Background

In recent years, research on the plant-specific AT-HOOK MOTIF NUCLEAR LOCALIZED (AHL) protein family has revealed their diverse roles in key developmental processes, including hypocotyl elongation, root architecture, somatic embryogenesis, vegetative phase change, flowering time, and longevity (Karami et al., 2021, 2020; Rahimi et al., 2022a; Street et al., 2008; Zhou et al., 2013). The *AHL* gene family is conserved across all land plants, with family sizes ranging from 10 genes in *Physcomitrella patens* (moss), 29 in *Arabidopsis thaliana* (Arabidopsis), 20-26 in *Oryza sativa* (rice) to 47 in carrot (Kim et al., 2011; Kumar et al., 2023; Machaj and Grzebelus, 2021; Zhao et al., 2014). Despite its considerable size and conservation, functional characterisation of *AHL* family genes has only recently begun and much remains to be uncovered regarding their roles in development and gene regulation.

AHL proteins bind to the minor groove of AT-rich DNA via a conserved AT-hook motif, which can also be found in other DNA-binding proteins, such as the High-Mobility Group A (HMGA) family (Fujimoto et al., 2004; Klosterman and Hadwiger, 2002). Based on domain composition, the AHL family is subdivided into two clades: clade A members contain a single, type A AT-hook domain, whereas clade B members possess a type B AT-hook domain, sometimes accompanied by an additional type A AT-hook domain (Zhao et al., 2014, 2013). In the nucleus, AHL proteins are proposed to form heterotrimeric complexes and to recruit other nuclear proteins through their PPC (Plant and Prokaryote Conserved) domains (Zhao et al., 2013).

While single *ahl* mutants often show very mild phenotypes, such as a slightly accelerated flowering time (Karami et al., 2020; Rahimi et al., 2022a), more striking phenotypic defects emerge when one of the characteristic AHL domains are disrupted. Deletion of the AT-hook domain (ΔAT-hook) abolishes DNA binding, while deletion of a six amino acid motif in the PPC domain (ΔG) or disruption of the PPC function via fusion to β-glucuronidase (GUS) impairs protein recruitment (Karami et al., 2021; Lin et al., 2007; Zhao et al., 2013). Such domain-specific mutations produce dominant-negative AHL variants that strongly interfere with endogenous AHL function. For example, expression of AHL15-ΔG or AHL15-GUS in a *ahl15/+* heterozygous mutant background inhibits both zygotic and somatic embryogenesis (Karami et al., 2021). Similarly, the *sob3-6* mutant, representing a *ΔAT-hook* allele of *AHL29*, displays a significantly elongated hypocotyl compared to wild-type and *ahl29* loss-of-function plants (Zhao et al., 2013). These findings indicate that AHL proteins function as part of multiprotein complexes with a high degree of functional redundancy. Loss of an AHL can be compensated by other family members, but incorporation of a dysfunctional AHL protein disrupts complex formation and function.

Due to the high functional redundancy between *AHL* genes, most studies have focused on phenotypes resulting from *AHL* overexpression, which are generally similar across different family members. For instance, overexpression of *AHL29/SOB3, AHL27/ESCAROLA/ORE7* or *AHL22* suppresses hypocotyl elongation (Street et al., 2008; Xiao et al., 2009), while overexpression of *AHL15, AHL19, AHL20, AHL22, AHL27,* or *AHL29* significantly delays flowering and enhances plant longevity (Karami et al., 2020; Street et al., 2008; Tayengwa et al., 2020; Yun et al., 2012). The delayed flowering phenotype is at least partially attributable to repression of *FT* expression, indicating that AHL proteins can function as transcriptional repressors (Tayengwa et al., 2020; Yun et al., 2012).

*AHL* overexpression has been associated with alterations in chromatin architecture. Lim et al. (2007) showed that *AHL27* overexpression reduces chromatin condensation, while Karami et al. (2021) demonstrated that *AHL15* overexpression decreases the number of nuclear foci and disrupts sister chromatid segregation during somatic embryogenesis, resulting in polyploid embryos. Several studies have also indicated that *AHL* overexpression is linked to histone modifications. Specifically, overexpression of *AHL* genes reduces HISTON 3 acetylation (H3Ac) and increases methylation of H3 at lysine 9 (H3K9me2) at AHL-bound loci, suggesting a role in promoting repressive chromatin states (Ng et al., 2009; Xiong et al., 2020; Xu et al., 2013; Yun et al., 2012). Consistent with this, AHLs have been shown to interact with a range of chromatin-modifying proteins, among which several histone deacetylases (HDA) and subunits of the SWR1 complex that exchanges H2A for H2A.Z (Lee and Seo, 2017; Xu et al., 2024, 2013; Yun et al., 2012). Notably, AHL22 was recently shown to interact with the transcriptional repressors FRS7 and FRS12 at matrix attachment regions (MARs) and to recruit HDA15 to the DNA (Xu et al., 2024).

Transcriptome analyses further support the role of AHLs in gene repression. RNA-seq data indicate that overexpression of AHLs leads to widespread transcriptional changes, with most differentially expressed genes being downregulated (Favero et al., 2020; Karami et al., 2022; Xu et al., 2024). Collectively, these findings demonstrate that *AHL* overexpression impacts chromatin structure, epigenetic regulation, and global gene expression, ultimately leading to pronounced effects on plant development. However, the molecular mechanisms by which AHL proteins are recruited to their genomic targets, how they influence histone modifications, and their broader role in chromatin architecture remain poorly understood.

The clade A AHL protein AHL15 has been shown to delay, and even reverse, developmental phase transitions, thereby extending plant longevity. *AHL15* overexpression induces somatic embryogenesis, whereas *ahl15* loss-of-function prevents BABY BOOM (BBM)-induced somatic embryogenesis. Together with its established role in zygotic embryogenesis, these findings indicate that AHL15 acts downstream of BBM during both zygotic and somatic embryogenesis (Karami et al., 2021). Furthermore, *ahl15* loss-of-function mutants exhibit accelerated vegetative phase change and early flowering, whereas both developmental transitions are delayed in *AHL15*-overexpressing plants (Rahimi et al, 2022a). Notably, *AHL15* overexpression converts herbaceous, monocarpic (short-lived) Arabidopsis plants into a woody, polycarpic (long-lived) plants phenocopying the *soc1 ful* double mutant. Conversely, the woody, polycarpic phenotype of *soc1 ful* double mutant is largely abolished in the *soc1 ful ahl15* triple mutant. Consistent with this genetic interaction, SOC1 and FUL directly bind the *AHL15* locus to repress its expression, demonstrating that AHL15 functions downstream of these two MADS-box transcription factors as a key regulator of plant longevity and woodiness (Karami et al., 2021; Rahimi et al., 2022b). In addition, we recently showed that leaf senescence is delayed by *AHL15* overexpression but significantly accelerated in the *ahl15* loss-of-function mutant (Luden et al., 2025), indicating that AHL15 promotes not only plant longevity but also organ longevity.

Despite its involvement in diverse developmental processes related to ageing, no direct genomic targets of AHL15 have yet been identified. Here, we combined RNA-seq analysis with Chromatin Immuno-Precipitation and Assay for Transposase-Accessible Chromatin sequencing (ChIP-seq and ATAC-Seq) to identify the genomic DNA binding sites of AHL15 in native- and overexpression conditions and investigate the effect of AHL15 on gene expression, chromatin accessibility, and its colocalization with several chromatin modifications. We show that a large set of loci is only bound by AHL15 upon overexpression, and that this coincides predominantly with downregulation of gene expression. In addition, we show that AHL15 does not affect chromatin accessibility, but that it preferentially binds at the boundaries of most epigenetic marks. Finally, we show that AHL15 binding sites largely overlap with those bound by GH1-GMGA2, an AT-hook protein shown to regulate chromatin architecture and gene loop formation (Zhao et al., 2021), implying that AHL15 could play a similar role.

## Results

### AHL15 binds near the TSS and TES of genes

To identify how AHL15 regulates juvenile plant development, we performed ChIP-seq experiments using young Arabidopsis plants expressing 3xFLAG-tagged AHL15. Because wild-type and *AHL15* overexpressing plants show significant phenotypic differences, we expressed the 3xFLAG-AHL15 fusion protein both under the native *(pAHL15:3xFLAG-AHL15* in *ahl15)* or a constitutively overexpressing (*p35S:3xFLAG-AHL15* in Col-0) promoter and pulled down AHL15-bound DNA from both lines (Figure 1A). In wild-type Arabidopsis, *AHL15* is expressed only in juvenile plants and its expression decreases significantly past the seedling stage (Rahimi et al., 2022a). We therefore performed ChIP on 10 day-old *pAHL15:3xFLAG-AHL15 ahl15* seedlings in which the *AHL15* promoter is still strongly active to identify the “native” DNA binding sites of AHL15. In contrast, *p35S:3xFLAG-AHL15* plants showed a stunted growth phenotype and a severe delay in flowering time (Additional file 1: Figure S1). To map the “overexpression” AHL15 targets, we performed ChIP-seq on 31-day old plants of this line before the onset of flowering.

**Figure 1:**
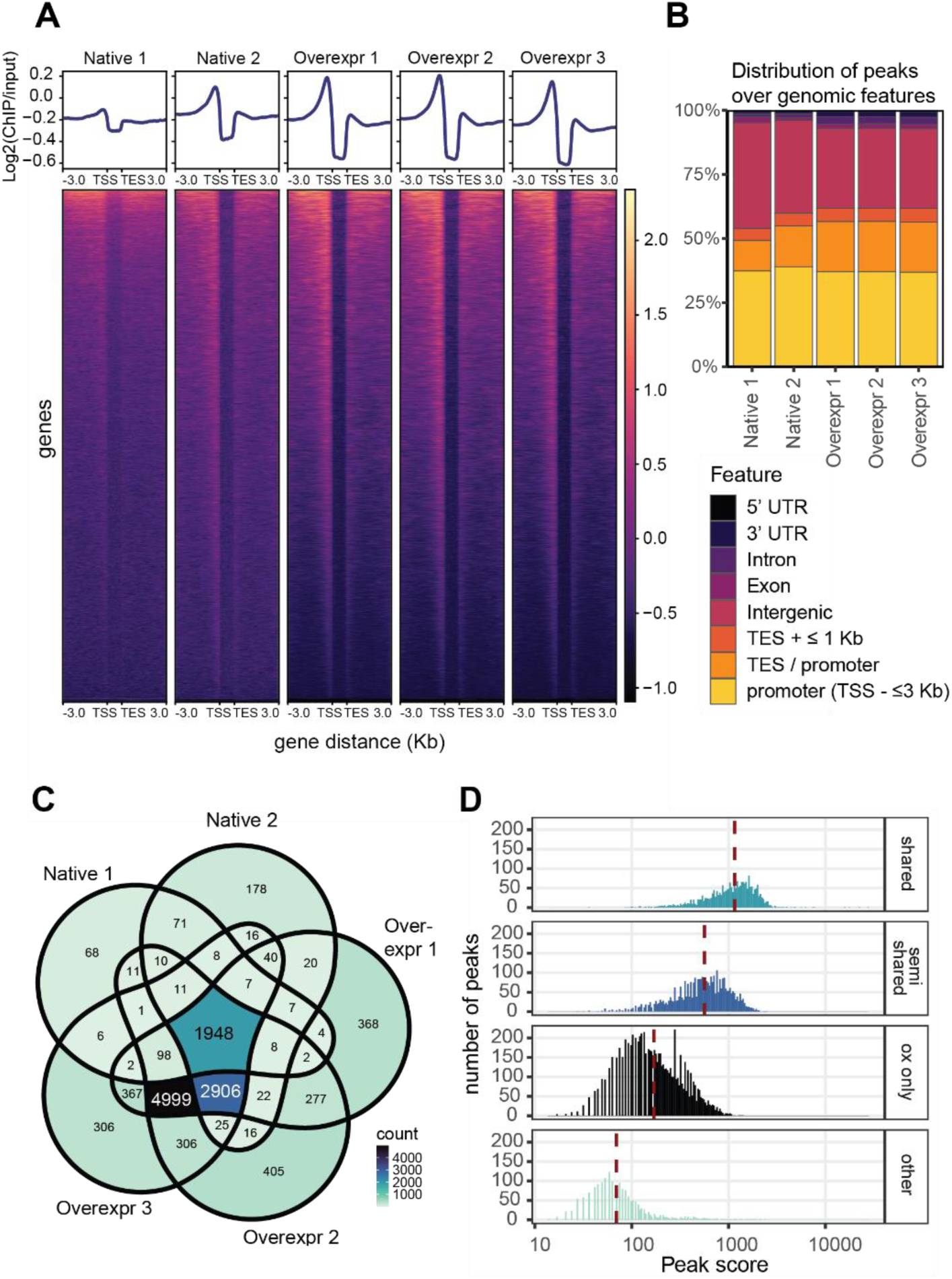
ChIP-seq of 3xFLAG-AHL15 in native and overexpression conditions. **A:** Heatmap and profile plots of AHL15 ChIP-seq peaks scaled over genes showing that AHL15 is enriched near the TSS and TES, and is largely absent over gene bodies. Shoots of ten day-old *pAHL15:3xFLAG-AHL15* were used for ChIP for the “native” condition, and 31 day-old *p35S:3xFLAG-AHL15* shoots were used for the “overexpression” condition. Profile plots show the log2 of ChIP reads over input reads. **B:** Annotation of 3xFLAG-AHL15 ChIP-seq peaks in **A** over genomic features of the Araport11 annotation of the Arabidopsis genome. TES: region up to 1 kb upstream of the TES. Promoter: region up to 3 kb downstream of the TSS. TES / promoter: regions where promoters overlap with the TES region of neighbouring genes. Intergenic: regions not covered by any of the before features. **C:** Venn diagram showing the overlap in peak annotations between the five ChIP-seq samples in **A**. 1948 peaks were identified in all five datasets (*shared*), 2906 peaks were shared between the three overexpression conditions and the second native ChIP (*semi-shared*), and 4999 peaks were identified in all three overexpression (OX) conditions (*OX only*). **D:** Histogram of the peak score of significant ChIP-seq peaks in *p35S:3xFLAG-AHL15* sample 2 (OX 2). Peaks identified in all five conditions have the highest peak score, followed by semi-shared peaks. OX-only peaks have a lower peak score than shared or semi-shared peaks, and peaks shared in less than 3 conditions have the lowest overall peak score. Reads were aligned with Bowtie2 and peak calling was performed with MACS2.

Our ChIP-seq data shows that AHL15 binds DNA abundantly in both native and overexpression conditions (Figure 1A). Under native conditions, we identified 2425 significant peaks that are shared between both replicates (Additional file 2). In the overexpression conditions, we identified over 14,000 significant peaks per replicate, of which 10,056 were shared between all three replicates, indicating that AHL15 binds DNA more broadly when overexpressed than in wild-type conditions, and that some DNA binding sites are unique to the overexpression condition (Additional file 2). Next, we mapped the AHL15 peaks over all genes in the Arabidopsis genome, which showed that AHL15 predominantly binds within 1 kb upstream of the transcription start site (TSS) of genes, and to a lesser extent near the transcription end site (TES) and is largely absent from gene bodies (Figure 1A). The peak of AHL15 occupancy in these regions was more pronounced in *AHL15* overexpression samples compared to the native condition, showing that AHL15 has a preference for the proximal promoter of genes regardless of whether these are genuine native targets or not. Annotation of the significant AHL15 ChIP-seq peaks over genomic features confirmed that the majority of AHL15 binding sites are located in promoters (up to 3 kb upstream of the TSS) or in intergenic regions (more than 3 kb upstream of the TSS or more than 1 kb downstream of the TES; Figure 1B). In addition, a substantial percentage of AHL15 ChIP-seq peaks was located in within 1 kb downstream of the TES or in regions where the promoter and TES of two genes overlapped (TES/promoter; Figure 1B). In overexpression conditions, the number of AHL15 ChIP-seq peaks mapping to intergenic regions was reduced in favour of promoter- and TES regions (Figure 1B), illustrating the increased binding of AHL15 to TSS and TES regions, as observed in Figure 1A.

To understand what loci are native targets of AHL15 and what loci are unique to the overexpression condition, we searched for overlap in significant ChIP peaks between samples and identified a set of 1948 peaks that are shared between all five ChIP-seq datasets (“shared”; Figure 1C). This set includes the promoters of genes such as *CYTOKININ OXIDASE3 (CKX3), WUSCHEL-RELATED HOMEOBOX 4 (WOX4),* and *TWIN SISTER OF FT (TSF)* (Additional file 2). In addition, 2906 peaks were shared between the second replicate of the native ChIP, which had a higher sequencing depth, and the three overexpression ChIPs (“semi-shared”), which included promoters of *SQUAMOSA PROMOTER BINDING-LIKE 9 (SPL9)* and *BRANCHED 1 (BRC1)*. We identified 4999 peaks that were common among all three overexpression datasets (“overexpression-only”) and included the promoters of *FLOWERING LOCUST (FT), SWEET10,* and *FLOWERING LOCUS C (FLC)* (Additional file 2). When comparing the distribution of peak scores in each category, it was clear that peaks that are shared by all datasets had a higher average peak score than those that are found only in overexpression datasets, indicating that AHL15 binds its native targets more strongly than its ectopic targets (Figure 1D). Thus, these data show that AHL15 can associate with DNA outside of its genuine native binding sites when overexpressed, but that the affinity for these regions is lower than for its native targets.

### AHL15 non-specifically binds AT-rich DNA sequences

Because AHLs are known to bind AT-rich DNA through their AT-Hook domain (Aravind, 1998; Franco-Zorrilla et al., 2014) and AHL15 binds a large number of promoter regions in both the native and overexpression conditions, we used the motif enrichment analysis tool HOMER (Heinz et al., 2010) to identify possible AHL15 binding motifs in the different datasets: shared, semi-shared, and overexpression-only AHL15 peaks as well as all significant peaks of the second overexpression replicate, which had the highest read depth (Figure 2A, Additional file 1: Figure S2-S4, Additional file 3). Although several AT-rich motifs were found to be significantly enriched in the different datasets, closer inspection showed that the frequency at which these motifs occurred within AHL15 peaks was in most cases less than 1%, making it unlikely to represent a true binding motif (Figure 2A, Additional file 1: Figure S2-S4). We argued that a true DNA binding motif would be present in a large portion of the AHL15 peaks (targets) and would be rare in other regions of the genome (background). In other words, the motif should have a high target : background ratio and be present in a high percentage of targets. Plotting the target : background ratio (Figure 2B) and the Log2 of the percentage of targets (Figure 2C) for all motifs identified by HOMER in the shared AHL15 peaks showed that the motifs having a high target : background ratio occurred in less than 1% of the AHL15 peaks, whereas motifs that were present in more than 1% of the AHL15 peaks had a low target : background ratio. Similar results were obtained for each of the AHL15 peak datasets (semi-shared, overexpression-only, and all AHL15 ox rep2 peaks), or when using only the 1000 shared peaks with the highest score (Additional file 1: Figure S5), and no motifs were shared between the four different datasets (Additional file 1: Figure S2-S4). Previous *in vitro* motif discovery experiments suggested several AT-rich sequences as binding sites for AHL proteins (Franco-Zorrilla et al., 2014; Fujimoto et al., 2004), which was confirmed by EMSA for AHL13 (Liu et al., 2026) and from motif enrichment analysis of AHL29 ChIP-seq peaks (Additional file 1: Figure S5, Favero et al., 2020). However, our results indicate that AHL15 does not recognize a *specific* DNA motif but is likely to identify its AT-rich target DNA based on other traits.

**Figure 2:**
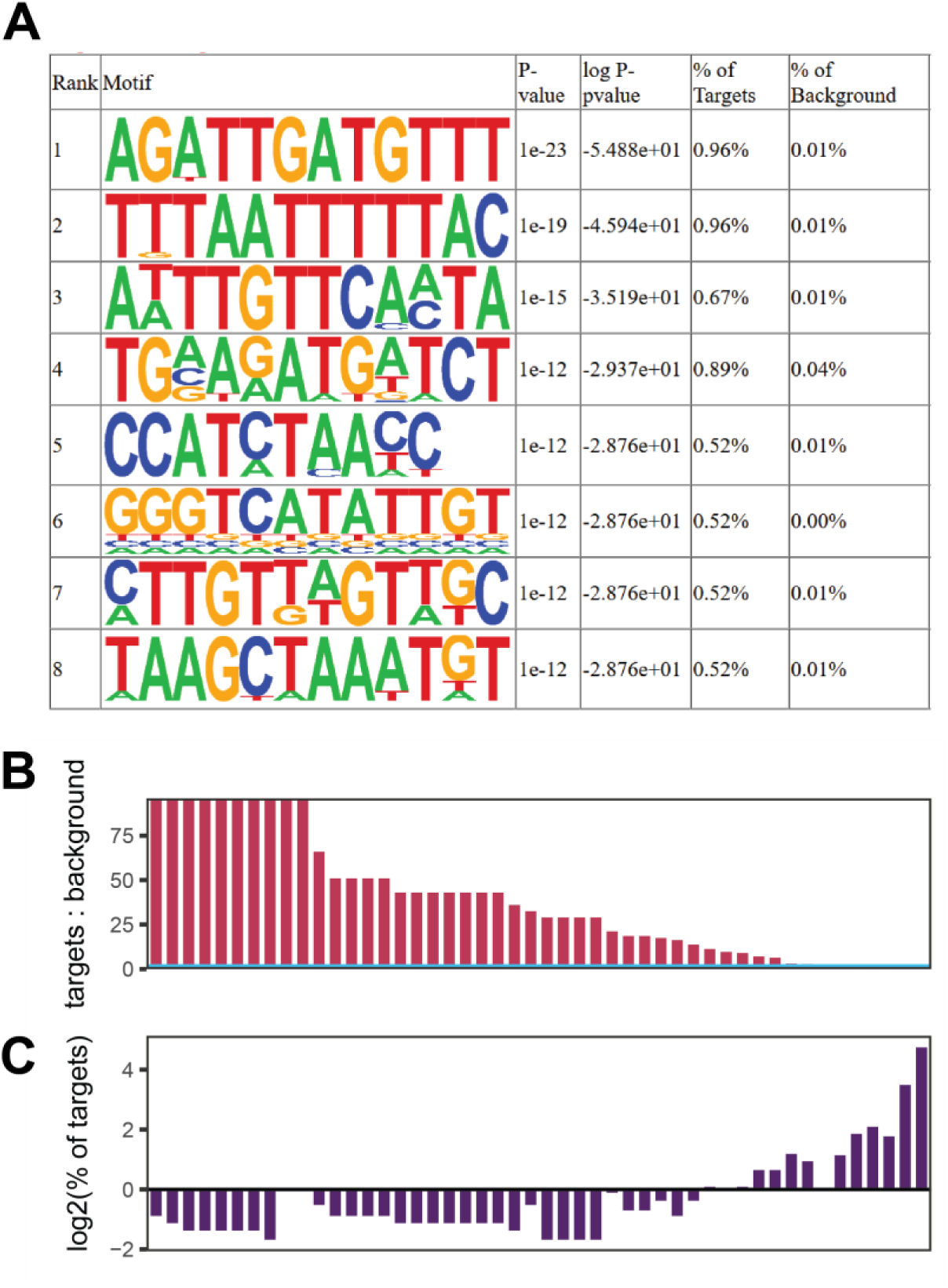
AHL15 binds to AT-rich DNA but does not recognize a specific DNA binding motif. **A:** Table showing the top 8 enriched motifs identified by HOMER2 within the set of shared peaks, with the p-value of motif enrichment, log_10_ of the p-value of motif enrichment, the percent of AHL15 peaks (targets) containing the motif, and the percent of background sequences containing the motif. **B:** Graph showing the AHL15 targets over background, meaning the number of times each motif (48 in total) is identified by HOMER2 in shared AHL15 peaks divided by its number over the entire genome (expressed as %; 100% means the motif is almost only present in AHL15 peaks). **C:** Graph showing the frequency with which each motif in **B** identified by HOMER2 occurs in shared AHL15 peaks, expressed as the log^2^ value of the % of shared AHL15 targets containing the specific motif (occurrence of more than 1% gives a positive value, 1% gives a value of 0, and less than 1% results in a negative value). Motifs in figures **B** and **C** are arranged by descending target : background value.

### Gene expression is predominantly repressed by AHL15

The large number of binding sites we identified in the native and overexpression conditions, as well as the strong phenotypic changes caused by *AHL15* overexpression suggest that AHL15 may directly or indirectly regulate the expression of many genes.

To determine genes whose expression is affected by *AHL15* overexpression and may underlie the developmental phenotypes associated with AHL15 activity, we used the Arabidopsis *p35S:AHL15-GR* line (Karami et al., 2020). In this line, a fusion protein of AHL15 and the glucocorticoid receptor (AHL15-GR) is constitutively overexpressed. Upon treatment with the steroid hormone dexamethasone (DEX), the fusion protein translocates from the cytosol to the nucleus, resulting in longevity-associated phenotypes similar to those observed in *p35S::AHL15* plants. These include delayed vegetative phase change and flowering (Figure 4C,D, Rahimi et al. (2022a)), prolonged vegetative development through delayed axillary meristem maturation, resulting in aerial rosette formation and polycarpy (Karami et al., 2020), enhanced secondary growth leading to woody stems (Rahimi et al., 2022b) and delayed leaf senescence (Luden et al., 2025). Ten day-old wild-type (Col-0) or *p35S:AHL15-GR* seedlings were submerged in 20 μM DEX or a mock solution for 15 minutes after which the solution was removed. Eight hours later seedling shoots were harvested for mRNA sequencing. Principal component analysis (PCA) of the RNA-seq data showed that the DEX-treated *p35S:AHL15-GR* samples grouped at one end, whereas the mock- or DEX-treated Col-0 or the mock-treated *p35S:AHL15-GR* samples grouped at the other end of the PCA plot, showing that DEX treatment on wild-type seedlings or submergence had a minimal effect on gene expression compared to DEX treatment on *p35S:AHL15-GR* plants (Additional file 1: Figure S7).

Induction of AHL15-GR activity by DEX treatment resulted in a substantial number of differentially expressed genes, including 1658 that were downregulated over twofold and 497 that were upregulated over twofold. Similar to other *AHL* genes, *AHL15* overexpression resulted a net downregulation of gene expression (Figure 3A, Additional file 4). Of the differentially expressed genes, 101 downregulated and 65 upregulated genes were also native targets of AHL15, and 501 downregulated and 155 upregulated genes were overexpression-only targets of AHL15 (Figure 3B-D, Additional file 5). Interestingly, the flowering-promoting genes *TSF* and *FT* and the *BRC1* and *BRC2* genes repressing axillary bud outgrowth were among the downregulated targets (Figure 3E), consistent with the delayed flowering and enhanced lateral branching phenotypes of *AHL15* overexpressing plants (Fletcher et al., 1999; Ji et al., 2010; Karami et al., 2020; Rahimi et al., 2022b).

**Figure 3:**
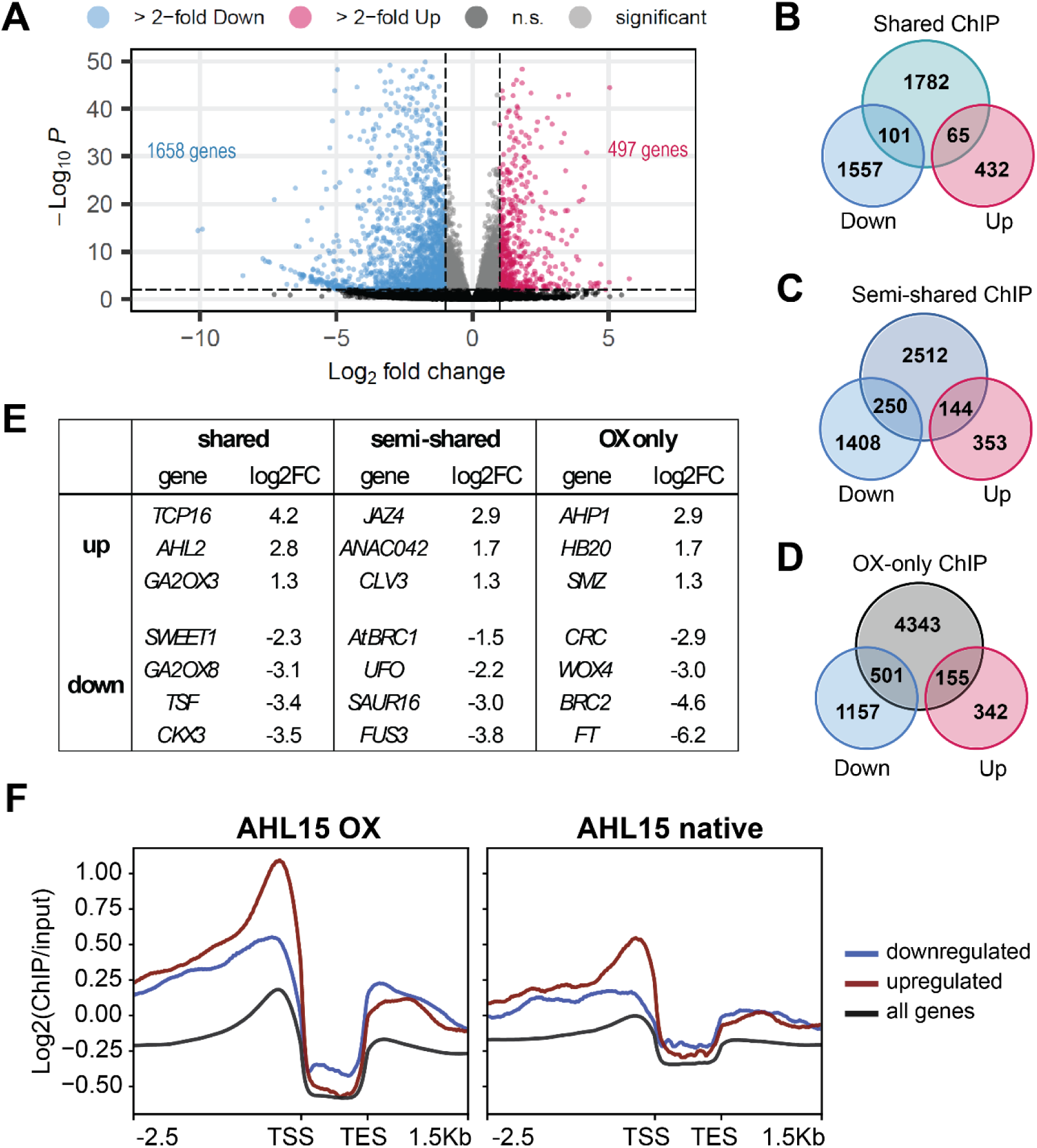
AHL15 binding at DNA regions near TES and TSS is enhanced for genes differentially expressed by AHL15-GR activation. **A:** Volcano plot showing differentially expressed genes in 10-day-old *p35S:AHL15-GR* seedlings after 8 hours treatment with DEX compared to the three control conditions (*p35S:AHL15-GR* seedlings treated with mock solution and Col-0 seedlings treated with either DEX or mock solution). Blue dots indicate genes that are significantly more than 2-fold downregulated, red dots indicate genes that are significantly more than 2-fold upregulated, light grey dots show genes that are significantly differentially expressed with a fold change of less than 2, and dark grey dots (black when overlapping) represent genes whose expression was not significantly altered. **B-D:** Venn diagrams showing overlap of differentially expressed genes with shared ChIP-seq peaks (**B**), semi-shared peaks (**C**), or overexpression-only peaks (**D**). **E:** Table showing a selection of genes shared between ChIP-seq and RNA-seq datasets identified in **B-D**. Log2FC: log^2^ fold change in gene expression. **F:** Profile plots of AHL15 ChIP-seq peaks over AHL15-bound differentially expressed genes vs all genes. AHL15 enrichment is stronger near the TSS of AHL15-bound differentially expressed genes and especially upregulated genes compared to the average of all genes. Near the TES, AHL15 enrichment is highest for downregulated AHL15-bound genes.

Knowing which genes are differentially expressed upon DEX treatment, we investigated the enrichment of AHL15 ChIP-seq peaks over the differentially expressed genes. To do this, we plotted the average enrichment of the native or overexpression AHL15 ChIP-seq datasets over AHL15-bound upregulated genes, AHL15-bound downregulated genes, or all genes. This revealed that AHL15 binding was stronger around the TSS and TES of AHL15-bound up- and downregulated genes compared to all genes (Figure 3F). Although the majority of differentially expressed genes was downregulated by DEX treatment, and downregulated genes represent a larger fraction of AHL15 DNA binding sites, we observed that AHL15 binding was highest around the TSS of upregulated genes, whereas AHL15 enrichment near theTES was slightly higher for AHL15-bound downregulated genes. In contrast, motif enrichment analysis of AHL15-bound differentially expressed genes again failed to reveal a specific DNA binding motif (Additional file 1: Figure S6). This suggests that strong AHL15 presence near the TSS of genes may lead to their upregulation, whereas strong AHL15 presence near both the TSS and the TES may lead to their downregulation.

### Chromatin accessibility is not directly affected by AHL15 induction

Previous research has shown that overexpression of *AHL15* or *AHL27* results in visible changes in chromatin structure (Karami et al., 2021; Lim et al., 2007). Additionally, our ChIP-seq data show that AHL15 binds to AT-rich DNA throughout the Arabidopsis genome with limited sequence specificity (Figure 2A, Additional file 1: Figure S2-S6). This suggests that AHL15 might alter chromatin accessibility globally. To test this, we performed ATAC-seq on 10-day-old *p35S:AHL15-GR* seedlings harvested at 30 minutes, 1 hour, and 8 hours after DEX or mock treatment. Surprisingly, we did not observe clear changes in chromatin accessibility at any of the time points: PCA showed that differences between replicates of the same condition were similar or larger than the differences between time points (Figure 4A). For example, ATAC-seq peaks at and around the *FT* locus were similar for all time points (Figure 4B), despite a strong downregulation of this gene upon DEX induction (Figure 3E). In addition, we did not detect any differentially accessible regions when comparing DEX- and mock treated samples at the 30m and 1h time points, and identified only 10 regions with differential accessibility for the 8h time point that did not correspond to differentially expressed genes (Additional file 6). Although we did not observe clear changes in chromatin accessibility at 8 hours after DEX treatment, the seedlings showed clear phenotypic changes at 2 weeks after DEX-treatment, resembling the phenotype of *p35S:AHL15* plants (Figure 4C; Additional file 1: Figure S1). In addition to a delay in bolting time of about 10 days in DEX-treated plants (Figure 4D), new leaves produced at the time of DEX treatment remained very small and did not develop to a full size, resulting in a severe reduction of the plant size (Figure 4C). Together with the large number of differentially expressed genes upon DEX treatment, these severe phenotypic changes indicate that AHL15 activation by DEX induces developmental reprogramming. However, our ATAC-seq data indicate that AHL15 does not affect chromatin accessibility directly, and that altered chromatin accessibility cannot explain the differential gene expression patterns observed at the 8h time point and the subsequent developmental reprogramming.

**Figure 4:**
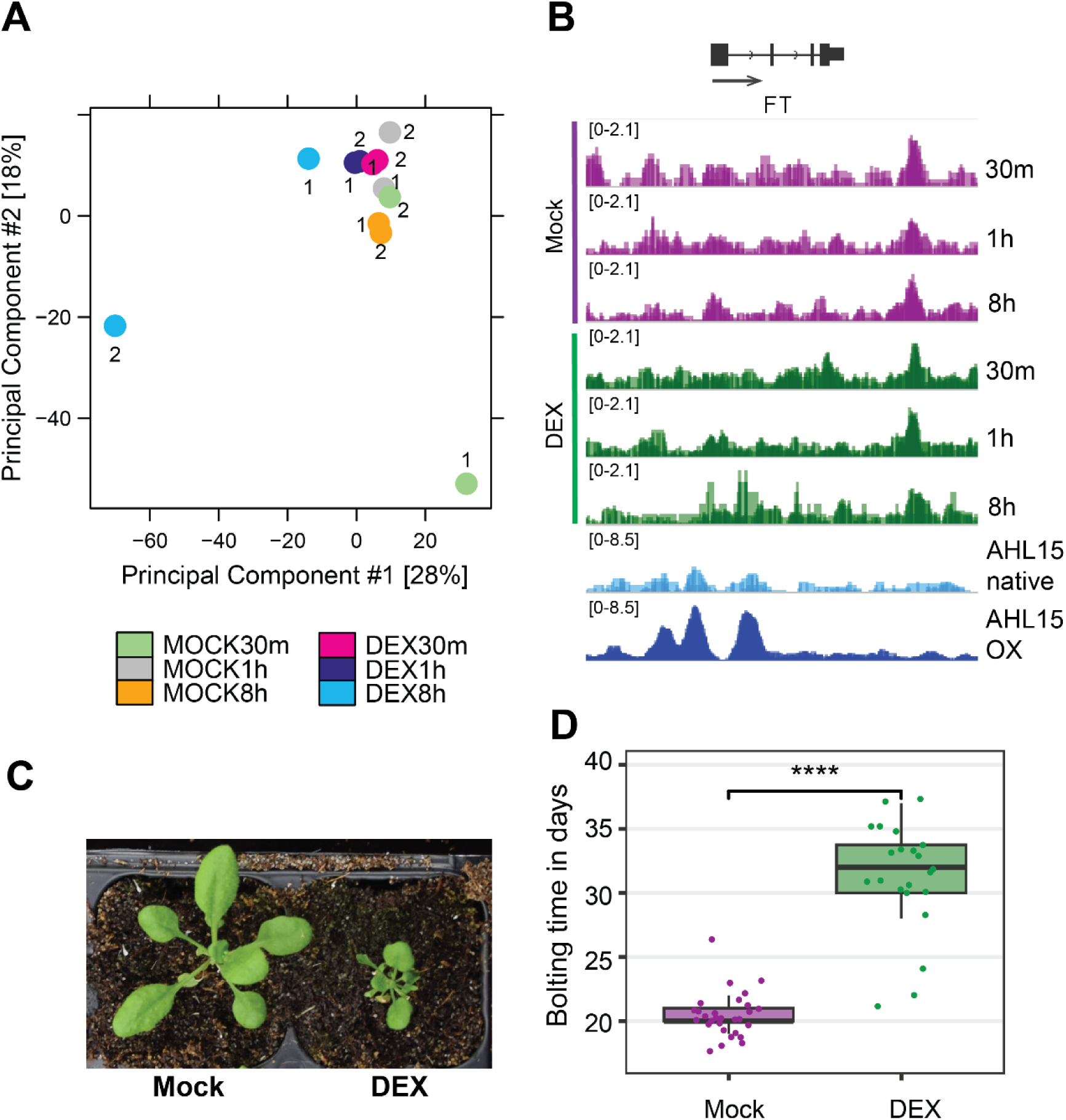
AHL15 delays plant developmental phase transitions in a chromatin accessibility-independent manner. **A, B:** ATAC-seq on 10 day-old *p35S:AHL15-GR* seedlings that were mock- or DEX treated for 30 minutes, 1 hour or 8 hours. **A:** Principal Component Analysis (PCA) of ATAC-seq samples, showing that most samples cluster together and that differences between replicates of the same condition (indicated by 1 and 2 for the same colour) are similar or larger than differences between conditions. **B:** ATAC-seq profiles (purple and green) and AHL15 ChIP-seq profiles (blue) at the *FT* locus, a gene that was strongly repressed 8 hours after DEX treatment. Each track shows the overlay of bigwig files from two (ATAC-seq and AHL15 native ChIP-seq) or three (AHL15 OX ChIP-seq) biological replicates. Arrows in the intron of the *FT* gene model indicate transcriptional direction. **C:** Phenotype of mock-treated and DEX-treated *p35S:AHL15-GR* plants at two weeks after DEX treatment (24 day-old plants). **D:** Bolting time of *p35S:AHL15-GR* plants following mock or DEX treatment, as shown in **C**. Seedlings were transferred to soil two days after DEX treatment and monitored for flowering time (n = 25; Wilcoxon test, p = 3.71·10^−9^). Plants were grown in long day conditions (16 hours light : 8 hours dark).

### AHL15 binds in regions with reduced epigenetic signatures

Because AHL15 does not have a measurable effect on chromatin accessibility as determined by ATAC-seq, we investigated whether AHL15 binding coincides with epigenetic markers. For this we used publicly available ChIP-seq data of several epigenetic marks in 10 day-old wild-type seedlings (Bourguet et al., 2021). We compared the distribution of these epigenetic markers as well as our ATAC accessibility data with that of AHL15 binding in the native and overexpression conditions by plotting the average enrichment of these marks over all Arabidopsis genes (Figure 5). This revealed that AHL15 binding is enriched in regions where chromatin accessibility (based on ATAC seq) is relatively low, and that AHL15 presence is depleted in the highly accessible regions near the TSS and in the gene body (Figure 5A). Similarly, AHL15 showed opposite patterns of enrichment compared to the deposition of H1 (Figure 5B), H2A variants (Figure 5C), and H3K9me1, H3K9me2, and H3K27me1 (Figure 5D). This opposing pattern of enrichment is clearly illustrated when inspecting the sequencing data at individual loci, showing that AHL15 peaks occur at regions where most or all other marks are depleted (Figure 5E). For example, the *TCP16* gene is upregulated in DEX-treated *p35S:AHL15-GR* seedlings, and AHL15 shows clear peaks flanking this gene in both native- and overexpression conditions (Figure 5E). At regions where AHL15 peaks are visible, a clear reduction in ATAC peaks and H2A.Z, and to a lesser extent H2A, H2A.X, H1, H3K9me1, H3K9me2, and H3K27me1 deposition can be seen. A similar pattern could be observed for all epigenetic marks except H3K9me1, and to a lesser extent H3K27me3, around the AHL15 peak near the *EULS3* locus, whose expression was unaffected by DEX treatment (Figure 5E). These patterns are also visible when plotting the enrichment of epigenetic marks around AHL15 ChIP-seq peaks. This analysis shows that chromatin accessibility (based on ATAC-seq) is reduced around the centre of AHL15 ChIP-seq peaks, and this reduction is most clear around strong AHL15 ChIP-seq peaks that were identified in both the native and overexpression condition and less pronounced around overexpression-only AHL15 peaks (Additional file 1: Figure S8). H2A, H2A.X, H2A.Z, and H1 deposition is also depleted most strongly around shared AHL15 peaks (Additional file 1: Figure S9), and H3K27me1 and H3K9me2 are both depleted around AHL15 peaks, but less strongly at shared AHL15 peaks than at overexpression-only peaks. Finally, H3K9me1 shows a slight enrichment at shared and semi-shared AHL15 peaks, illustrated by the peak seen at the *EULS3* locus (Figures 5E, S7E). Taken together, these analyses clearly show that AHL15 binds at regions that are depleted of most epigenetic marks, which could be explained in different ways. First, it is possible that AHL15 is repelled by the presence of specific marks such as H2A.Z, and therefore binds only in regions where these marks are absent. Alternatively, it is possible that AHL15 has a preference for regions with reduced chromatin accessibility, which would explain the opposing patterns of AHL15 binding and ATAC-seq peaks. A third possibility is that AHL15 binding results in the removal of the deposition of epigenetic marks, but the depletion of most histone modifications around overexpression-specific AHL15 peaks does not support this idea.

**Figure 5:**
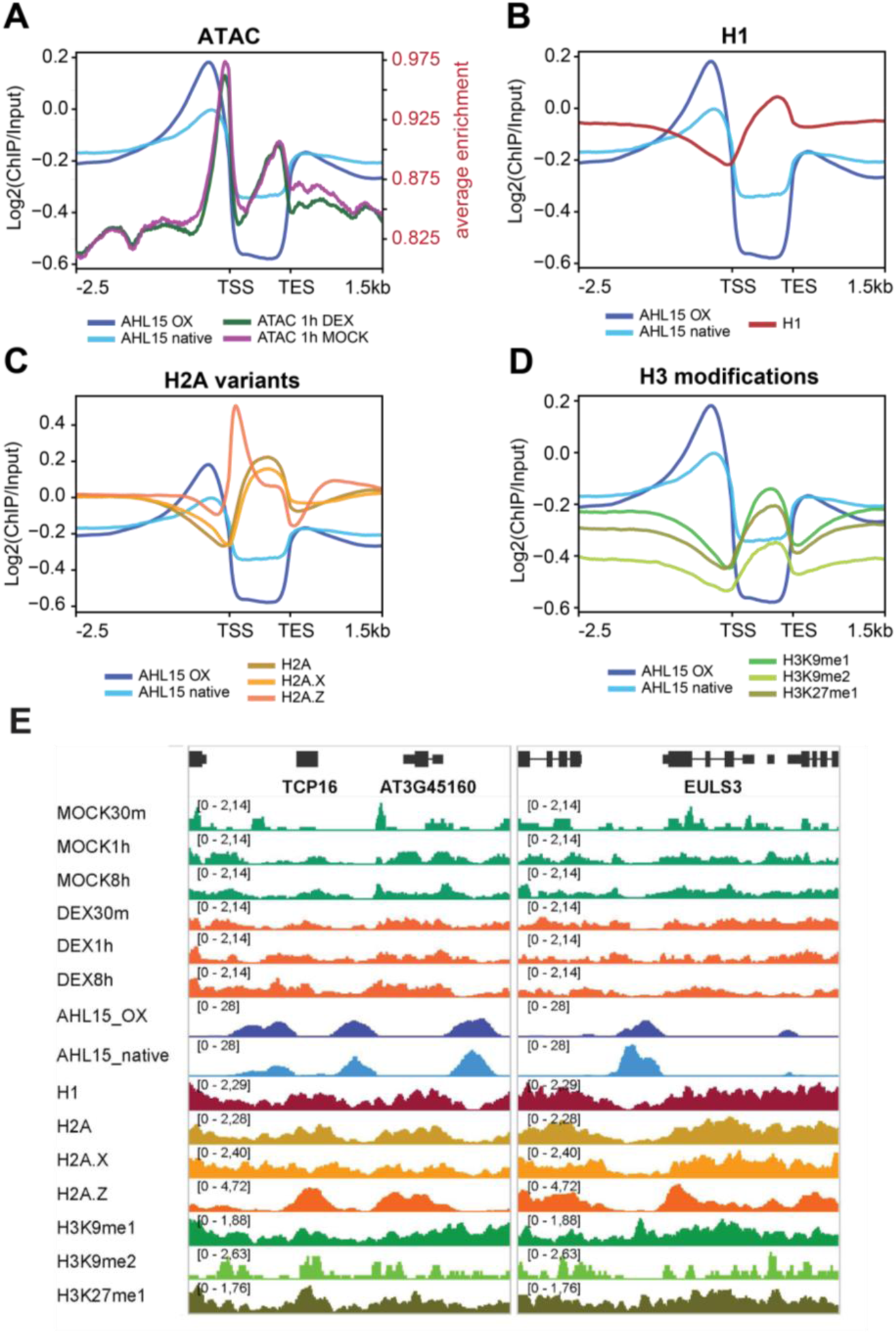
AHL15 binds DNA regions characterised by reduced chromatin accessibility and a depletion of epigenetic marks. **A-D:** ChIP-seq profiles of *pAHL15:3xFLAG-AHL15* (AHL15 native) and *p35S:3xFLAG-AHL15* (AHL15 OX) scaled over all Arabidopsis genes and compared to the ATAC-seq profile of *p35S:AHL15-GR* plants treated with DEX or mock solution for 1 hour (**A**), or to the ChIP-seq profiles of Histone 1 (H1) (**B**), Histone 2 variants H2A, H2A.X and H2A.Z (**C**), or the Histone 3 methylation marks H3K9me1, H3K9me2, and H3K27me1 (**D**). **E:** Alignment of the AHL15 native (AHL15_native) and overexpression (AHL15_OX) ChIP-seq profiles (light and dark blue, respectively) at the *TCP16* and *EULS3* loci with the averaged ATAC-seq profiles of ten day-old *p35S:AHL15-GR* seedlings that were mock-(green) or DEX (orange) treated for 30 minutes, 1 hour or 8 hours. Below the AHL15 ChIP-seq profiles, the ChIP-seq profiles of the epigenetic marks mentioned in **B-D** are shown.

### AHL15 binds genes at both ends and may be involved in gene looping

AHL15 binds predominantly near the TSS of genes and is also enriched near the TES (Figure 1A, B, Figure 5E), indicating that its presence at both ends of genes may be inherent to its mechanism of transcriptional regulation. Like AHL proteins, members of the globular H1 domain containing high-mobility group A (GH1-HMGA) protein family have AT-hook DNA-binding domains that bind the minor groove of AT-rich DNA and can affect local chromatin structure (Charbonnel et al., 2018; García-Heras et al., 2009; Ozturk et al., 2014). It was shown that GH1-HMGA proteins bind at both the 5’ and 3’ end of the *FLC* locus and thereby disrupt the formation of a gene loop associated with active transcription, resulting in a reduced transcription of *FLC* and an accelerated flowering time (Zhao et al., 2021). As AHL and GH1-HMGA proteins share the same type of DNA-binding domain, and because AHL15 peaks also flank genes at both ends, we compared our AHL15 ChIP-seq data to previously published ChIP-seq data of GH1-HMGA2/HON5. GH1-HMGA2/HON5 is enriched near the TSS and TES of genes in a similar pattern as AHL15, indicating that both proteins may bind at the same position (Figure 6A). To test this, we plotted the enrichment of GH1-HMGA2/HON5 ChIP-seq reads over AHL15 peaks, and found that GH1-HMGA2/HON5 reads are strongly enriched at these positions (Figure 6B). GH1-HMGA2/HON5 binding is strongest at canonical AHL15 binding sites, as it was most enriched at shared AHL15 peaks and weaker at overexpression-only peaks. Comparison of AHL15 and GH1-HMGA2/HON5 enrichment at specific loci further confirmed that they bind at the same position. For the self-looping gene *FLC*, which was not differentially expressed upon DEX activation of AHL15, the GH1-HMGA2/HON5 peak near the TSS of *FLC* was also present in the AHL15 overexpression datasets, as well as a weaker peak near the *FLC* TES (Figure 6C). Similarly, for the AHL15-upregulated self-looping gene *RCD1*, both GH1-HMGA2/HON5 and AHL15 bind at both ends of the gene (Figure 6C). By comparing the AHL15 ChIP-seq data with 1766 previously identified self-looping genes (Liu et al., 2016), we found that AHL15 binding at self-looping genes is significantly enhanced in overexpression conditions compared to native conditions (Figure 6D). In addition, at least 91 AHL15-bound self-looping genes were also upregulated, and 77 AHL15-bound self-looping genes were downregulated upon DEX treatment (Figure 6E, Additional file 7). No conserved DNA motif was discovered within the set of self-looping AHL15-bound genes, but this could be in part explained by the small size of sequences used for the analysis (Additional file 1: Table S1). Taken together, these comparisons show that, like GH1-HMGA proteins, AHL15 is likely to be involved in regulating gene loop formation and by this may affect the expression of at least some its target genes.

**Figure 6:**
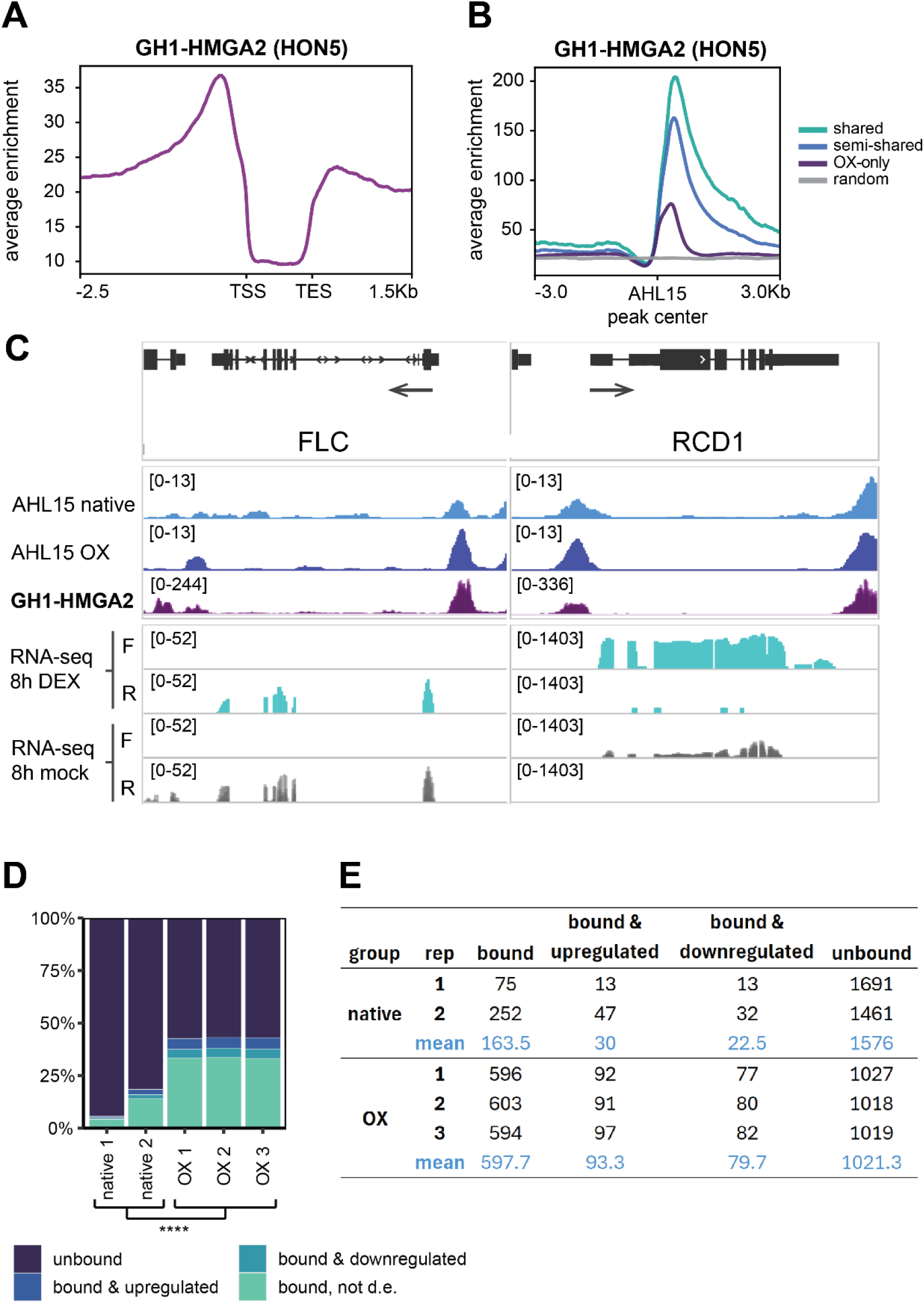
AHL15 shows overlapping DNA binding with GH1-HMGA2/HON5, suggesting a role in gene looping. **A:** Profile plot of GH1-HMGA2/HON5 ChIP-seq reads from (Zhao et al., 2021) scaled over Arabidopsis genes, showing a similar enrichment pattern as AHL15. **B:** GH1-HMGA2/HON5 ChIP-seq reads are were plotted over three classes of AHL15 ChIP-seq peaks presented in Figure 1C shared (identified in native and overexpression conditions, n = 1948), semi-shared (identified in one native sample and in all overexpression samples, n = 2906), and OX-only (identified in all three overexpression samples, n = 4999). **C:** AHL15 and GH1-HMGA2/HON5 ChIP-seq peaks around the *FLC* and *RCD1* loci. The RNA-seq tracks below indicate mapped reads per strand; the *FLC* gene is located on the reverse strand (R) and *RCD1* is located on the forward (F) strand. The ChIP-seq and RNA-seq tracks show an average of biological replicates of the same condition. **D:** Graph showing the percentages of self-looping genes in the individual native or overexpression (OX) ChIP-seq samples not bound by AHL15, bound by AHL15 and upregulated upon DEX treatment, bound by AHL15 and downregulated upon DEX treatment, and bound by AHL15 but not differentially expressed (d.e.). (**** indicates significant difference in the χ^2^ test, p < 0.0001). **E:** Table showing the number of self-looping genes identified in each ChIP-seq sample and sub-group shown in **D.**

## Discussion

Overexpression of various *AHL* genes- particularly *AHL15*-has been shown to induce developmental reprogramming, delaying or even reversing ageing-related developmental transitions such as vegetative phase change, the floral transition, and leaf senescence (Karami et al., 2021, 2020; Lim et al., 2007; Rahimi et al., 2022a; Street et al., 2008). However, the underlying molecular mechanisms driving these developmental changes have remained largely unclear. Here, we mapped the genome-wide DNA binding sites of AHL15 expressed under its own (native) or overexpressed from the constitutive *p35S* promoter, and show that when overexpressed, AHL15 is present throughout the genome at many ectopic binding sites, but binds more strongly at its native targets. AHL15 binds AT-rich DNA regions near the TSS and TES and is largely absent from gene bodies. Despite this clear positional preference, we did not identify a convincingly conserved binding motif, as the most frequently occurring motifs enriched in AHL15-bound regions also show high occurrence in the genomic background. This suggests that while showing a clear preference for AT-rich DNA, AHL15 and likely other AHL proteins do not recognize a specific DNA sequence but are recruited to their AT-rich target sites in another way.

Our experiments with the DEX-inducible *p35S:AHL15-GR* line show that *AHL15* overexpression causes strong transcriptional and phenotypic changes, with a majority of differentially expressed genes being downregulated. About one third of these differentially expressed genes was also bound by AHL15, indicating that AHL15 directly regulates the expression of a large number of genes. However, ATAC-seq in the same conditions showed that the changes in transcriptome induced by AHL15-GR activation do not coincide with changes in chromatin accessibility, suggesting that AHL15 regulates transcriptional activation or repression in an accessibility-independent manner. We also compared the positions of AHL15 peaks with ATAC-generated accessibility data, and saw that AHL15 binds regions with relatively low accessibility, just upstream of the highly accessible regions around the TSSs. As AHL15-GR activation by DEX treatment did not cause changes in the ATAC-seq profile within 8 hours, it is likely that AHL15 binding does not reduce chromatin accessibility itself, but is rather recruited to regions with reduced accessibility. However, it remains unclear what attracts AHL15 or other AHL proteins to a specific locus.

We also investigated whether AHL15 peaks overlap with heterochromatin-associated epigenetic marks such as H1, H3K9me1, and H3K9me2, and observed a clear pattern in which AHL15 flanked most of these marks. AHL proteins were initially discovered as components of the nuclear matrix, and AHL1 was shown to localize at the boundary between heterochromatic regions and H3K4me-marked regions (Fujimoto et al., 2004). Our data show that AHL15 similarly flanks other epigenetic signatures, as we saw no overlap of AHL15 with any mark except a weak overlap with H3K9me1 deposition. It is likely that AHL15 preferentially binds to regions devoid of epigenetic marks, but we cannot exclude the possibility that AHL15 induces removal of some histone variants or epigenetic modifications, for example that of H2A.Z or H1, which were both more strongly depleted around shared (native and OX) AHL15 peaks than around OX-only peaks. Previous studies have shown that overexpression of different clade A *AHLs* results in increased H3K9me2 and decreased H3Ac deposition at the *FT* and *FLC* loci (Ng et al., 2009; Xu et al., 2013; Yun et al., 2012), but these findings are likely to represent indirect effects of enhanced *AHL* expression rather than direct effects of AHL binding to the loci that were investigated. Genome-wide profiling of epigenetic marks in DEX-induced and non-induced AHL15-GR plants should give conclusive evidence of the effect of AHL15, and by proxy other AHLs, on the deposition of epigenetic marks.

Interestingly, AHL15 peaks overlap with those of GH1-HMGA2/HON5, a protein that was previously shown to interfere with gene loop formation between the 5’ and 3’ ends of the *FLC* locus, thereby hindering *FLC* transcription (Zhao et al., 2021). The similarity in DNA-binding domains between AHL15 and GH1-HMGA2/HON5 explains their overlap in DNA binding sites, and suggests that AHL15 may play a similar role as GH1-HMGA2/HON5 in regulating loop formation. HMGA-family proteins contain four AT-hook domains, whereas clade A AHLs have a single AT-hook domain (Klosterman and Hadwiger, 2002; Zhao et al., 2014). AHLs are known to form complexes with themselves and with other AHLs (Zhao et al., 2013), resulting in a complex that contains three AT-hooks, and this property could be necessary to bind AT-rich DNA with sufficient strength to affect its 3D confirmation. This could explain the strong effect of dominant-negative AHL alleles on the phenotype: the absence of an AT-hook domain in ΔAT-hook variants likely reduces the affinity for DNA of the entire AHL complex they are part of, thereby reducing the potential of entire complexes to bind DNA and modify its architecture. On the other hand, AHL complexes that incorporate the ΔG variant can still bind DNA, but have a reduced ability to recruit interacting proteins that could be required to alter the chromatin architecture. In both cases, the reduced ability of AHL complexes to affect DNA structure likely interferes with the proper regulation of gene expression, explaining the strong phenotypes observed in dominant-negative mutants (Karami et al., 2021, 2020; Zhao et al., 2013). In addition, AHL complexes and HMGAs might compete for the same regions in the DNA and have an effect on gene loop formation via association with different interaction partners. Although the expression of *FLC* was not affected in DEX-treated seedlings, several other self-looping genes were differentially expressed upon AHL15-GR activation. It would therefore be interesting to study whether these previously reported self-loops are altered upon AHL15-GR activation, whether AHL15 facilitates or disrupts these loops, and whether this results in altered transcription at these loci.

Previous research has shown that TFs that activate gene expression preferentially bind to regions near the TSS, whereas repressors are more likely to bind both near the TSS and near the TES (Franco-Zorrilla et al., 2014). In line with this, we observed that AHL15 enrichment was enhanced around the TSS of AHL15-bound upregulated genes, whereas enrichment near the TES was highest for AHL15-bound downregulated genes. This suggests that strong association of AHL15 with the TSS of genes relative to the TES enhances their expression and that a more equal distribution of AHL15 binding near the TSS and TES represses their expression. However, the effect of AHL15 binding at these regions on gene loop formation remains unclear.

Finally, our ATAC-seq data shows that AHL15 does not directly affect chromatin accessibility. This result is surprising, as earlier reports have shown that overexpression of *AHL15* or *AHL27* causes visible changes in the intensity of heterochromatin foci in the nucleus, suggesting a reduced chromatin compaction (Karami et al., 2021; Lim et al., 2007). It is possible that AHL15 induces chromatin opening on a slower time scale than measured here, but our finding that AHL15 may act similar to GH1-HMGA2/HON5 in the 3D organization of DNA could offer an alternative explanation for this phenomenon. GH1-HMGA2/HON5 was shown to disrupt chromatin looping on a local scale, but the authors did not investigate the effect of this protein on the global 3D structure of the DNA (Zhao et al., 2021). As chromatin architectural modifiers, it is likely that both GH1-HMGA and AHL proteins can affect chromatin looping on short-range as well as long-range scales. Here, we have shown that AHL15 binds in regions depleted in histone modifications and other epigenetic signatures, and similarly, AHL1 was shown to be present at the boundaries between H3K4me-marked euchromatin and DAPI-stained heterochromatin foci (Fujimoto et al., 2004). It is therefore possible that excessive AHL presence alters or disrupts these boundaries, causing changes in heterochromatin compartmentalization that result in the disappearance of heterochromatin foci that is observed in *AHL* overexpression lines (Karami et al., 2021; Lim et al., 2007).

## Conclusions

This study provides a genome-wide analysis of AHL15 binding sites in both native and overexpression conditions, revealing thousands of AHL15 peaks predominantly located at ca. 700 bp upstream of the TSS or near the TES of genes. Combined with transcriptomic data from plants with induced AHL15 activity, this identified a comprehensive set of direct AHL15 targets such as *FT, FUS3,* and *CKX3*, that explain the developmental reprogramming phenotypes observed in *AHL15* overexpressing plants. The surprising lack of changes in chromatin accessibility upon AHL15 induction, along with the depletion of chromatin accessibility and other epigenetic marks (H1, H2A variants, H3 modifications) at AHL15 binding sites provide evidence that AHL15 modulates transcription without modifying the epigenetic landscape. In line with this, the DNA binding profile of AHL15 strongly overlaps with that of the AT-hook containing GH1-HMGA2/HON5 protein that was previously shown to control gene expression by altering gene loop formation, suggesting a similar function for AHL15. Taken together, our data provide strong support for a role of AHL15 as a regulator of transcription via modification of chromatin architecture.

## Materials & methods

### Molecular cloning

To create the *p35S:3xFLAG-AHL15* vector, a *3xFLAG-AHL15* construct was synthesized at Baseclear (Leiden, the Netherlands) and inserted to the PMDC32 vector (Karimi et al., 2007) by digestion with SpeI and KpnI, and the hygromycin resistance gene was replaced by the NptII kanamycin resistance gene using XhoI restriction sites at both ends. For the *pAHL15:3xFLAG-AHL15* vector, the AHL15 promoter (up to 3 kB upstream of the *AHL15* TSS) was amplified by PCR with inclusion of the HindIII restriction site at the 5’ end and the KpnI and SacI sites at the 3’ end. Digestion with HindIII and SacI was used to replace the *35S* promoter in the PMDC32 vector with *pAHL15,* and the 3xFLAG-AHL15 construct was subsequently introduced by digestion with KpnI and SpeI. The final *p35S:3xFLAG-AHL15* and *pAHL15:3xFLAG-AHL15* vectors were introduced into *Agrobacterium tumefaciens* strain *AGL1* by electroporation (Dulk-Ras and Hooykaas, 1995).

### Plant material and growth conditions

The Arabidopsis *p35S:AHL15-GR* line and *ahl15* (SALK_040729) loss-of-function mutant have been described previously (Karami et al., 2020). *p35S:3xFLAG-AHL15* or *pAHL15:3xFLAG-AHL15* constructs were transformed to Arabidopsis by floral dip (Clough and Bent, 1998). The overexpression construct was expressed in the Col-0 background and *pAHL15:3xFLAG-AHL15* was expressed in the *ahl15* loss-of-function mutant background. One single locus line homozygous for the T-DNA insertion was chosen for each construct (from multiple independent transgenic lines) based on the stability of the transgene expression (in the case of *p35S:3xFLAG-AHL15)* or based on complementation of the *ahl15* phenotype (for *pAHL15:3xFLAG-AHL15 ahl15)* and used for experiments. Seeds were sterilized in 2% sodium hypochlorite for 2 minutes and washed with sterile milliQ water (mQ) before plating on ½ MS medium (Murashige and Skoog, 1962) with 1% sucrose and 0.7% agar (for horizontal growth) or 1.2% agar (for vertical growth) and then stratified for three days at 4 °C.

For native ChIP, RNA-seq and ATAC-seq, seeds were sown on ½ MS in square plates which were placed vertically in a growth chamber and grown for 10 days at 21 °C and a 16-hour photoperiod before treatment and/or harvesting. For native ChIP, *p:3xFLAG-AHL15 ahl15* seeds were sown on ½ MS with 20 mg/L hygromycin, and shoots were harvested for fixation after 10 days of vertical growth. For RNA-seq and ATAC-seq, *p35S:AHL15-GR* and Col-0 seeds were grown vertically on ½ MS without antibiotics until drenching and harvesting at 10 days.To check the effect of DEX treatment of the 10-day old *p35S:AHL15-GR* seedlings on later plant development, DEX or mock treated seedlings that were not harvested for RNA-seq or ATAC-seq were transferred to soil 2 days after treatment and grown at 21 °C, a 16-hour photoperiod and 70% relative humidity until all plants bolted. For overexpression ChIP, *p35S:3xFLAG-AHL15* seedlings were grown on ½ MS with 15 mg/L kanamycin for 10 days and then transferred to soil and grown in a growth chamber at 21 °C, a 16-hour photoperiod and 70% relative humidity for an additional 21 days prior to harvesting whole shoots for ChIP.

### ChIP-seq

The ChIP protocol used was adapted from the protocols by Kaufmann et al. (2010) and Chouaref et al. (2018). For each ChIP, 1 g of shoot tissue from 31 day-old *35S:3xFLAG-AHL15* plants or 10 day-old *pAHL15:3xFLAG-AHL15* seedlings was harvested and fixated for 10 minutes on ice and under vacuum in 1% formaldehyde in ice-cold MC buffer (10 mM sodium phosphate buffer pH 7, 50 mM NaCl, 0.1 M sucrose). The fixation reaction was stopped by saturation with glycine, and samples were washed in ice-cold MC buffer, dried with tissue paper, and stored at −80 °C. For nuclei isolation, frozen fixated tissue was ground to a very fine powder in liquid nitrogen using a mortar and pestle, and was then resuspended in 15 mL M1 buffer (10 mM sodium phosphate buffer pH 7, 0.1 M NaCl, 1 M 2-methyl 2,4-pentanediol, 10 mM β-mercaptoethanol) and gently shaken at 4 °C for 30 minutes. Suspensions were subsequently filtered through respectively a 100 µM and a 70 µM nylon mesh, both of which were washed with an additional 5 mL M1 buffer to extract remaining nuclei. The filtered suspension was then centrifuged for 20 minutes at 1000x*g* at 4°C, and pellets were resuspended in 20 mL M2 buffer (10 mM sodium phosphate buffer pH 7, 0.1 M NaCl, 1 M 2-methyl 2,4-pentanediol, 10 mM β-mercaptoethanol, 10 mM MgCl_2_, 0.5% Triton X-100). After 30 minutes of gentle shaking at 4 °C, the suspension was centrifuged for 10 minutes at 1000x*g* at 4 °C, the supernatant was discarded and the pellet was resuspended again in 5 mL M2 buffer. After 10 minutes of gentle shaking followed by 10 minutes centrifugation at 1000x*g* at 4 °C, the pellet was resuspended in 1 mL sonication buffer (10 mM sodium phosphate buffer pH 7, 0.1 M NaCl, 0.5% Sarkosyl, 10 mM EDTA) and aliquoted at 300 μL in 1.5 mL protein lobind tubes (Eppendorf, Germany). Samples were then sonicated in a Bioruptor sonicator (Diagenode, Seraing, Belgium) on HIGH for 5 rounds of 10 cycles of 30 seconds ON / 30 seconds OFF to a fragment size of ∼300 bp. After sonication, tubes of the same replicate were pooled and spun for 10 minutes at 12000x*g at* 4 °C, and the supernatant was transferred to a new tube and stored at −20 °C.

For the immunoprecipitation, 50 µg chromatin was used per IP (approximately 250 µL sonicated material), and 3.5 volumes IP buffer (50 mM HEPES pH 7.5, 150 mM NaCl, 5 mM MgCl_2_, 10 µM ZnSO_4_, 1% Triton X-100, 0.05% SDS) and 25 µL protein G dynabeads (Thermo Fischer, Lithuania) conjugated to mouse monoclonal Anti-FLAG M2 antibodies (F1804; Sigma-Aldrich, Darmstadt, Germany) were added and rotated at 4 °C overnight. After this, beads were washed once in 1 mL of each of the following buffers: IP buffer, high salt buffer (500 mM NaCl, 0.1% SDS, 1% Triton X-100, 2 mM EDTA, 20 mM Tris-HCl pH 8), LiCl buffer (10 mM Tris-HCl pH 8, 1 mM EDTA, 1% NP-40, 1% sodium deoxycholate, 0.25 M LiCl), and twice in TE buffer (10 mM Tris-HCl pH 8, 1 mM EDTA), and then resuspended in 50 µL TE buffer. Samples were de-crosslinked overnight at 65 °C after adding 450 μL of decrosslinking buffer (50 mM Tris-HCl pH 8, 10 mM EDTA pH 8, 250 mM NaCl, 1 % (w/v) SDS). DNA was cleaned by phenol: chloroform extraction, ethanol precipitated, and resuspended in milliQ and treated with RNAse A (Thermo Scientific, Vilnius, Lithuania) for 30 minutes at 37 °C, and then stored at −20 °C until sequencing. ChIP samples were sequenced with the Illumina Novaseq 6000 at Macrogen (Amsterdam, the Netherlands) with 20 million 150 bp paired-end reads.

### RNA-seq

Ten day-old seedlings (Col-0 and *p35S:AHL15-GR*) grown on plates with ½ MS medium (see above) were flooded at zeitgeber time 1 with 35 mL 20 μM DEX in milliQ or with a mock solution with an equivalent volume of DMSO for 15 minutes, after which the solution was siphoned off. The plates were returned to the growth chambers and five seedlings per replicate (n=3) of each condition (*p35S:AHL15-GR* + DEX, *p35S:AHL15-GR* + mock, Col-0 + DEX, Col-0 + mock) were harvested (shoot only) after eight hours and snap-frozen in liquid nitrogen. The frozen plant material was pulverized with a metal bead in a Tissuelyser (Qiagen, Hilden, Germany) and RNA was isolated using Trizol© (Thermo Fischer, Vilnius, Lithuania) with an additional ethanol wash step. For sequencing, 3 μg RNA was aliquoted and treated with DNAse-I (Thermo Fischer, Vilnius, Lithuania) for 30 minutes at 37 °C. Library preparation and sequencing with the Illumina Novaseq 6000 (150 bp paired-end, 10 million reads) was done at Baseclear (Leiden, the Netherlands).

### ATAC-seq

Ten day-old *p35S:AHL15-GR* seedlings grown in plates with ½ MS (see above) were treated with DEX or mock solution as for RNA-seq. For each replicate, approximately 50 DEX- or mock-treated seedlings were harvested (shoot only) at 30 minutes, 1 hour, or 8 hours after treatment, dried, and snap-frozen in liquid nitrogen and stored at −80 °C. Nuclei isolation was done by chopping frozen seedlings on ice in 2 mL nuclei isolation buffer (INGR) and subsequent filtering through a 40 μM mesh according to the protocol by Wang et al. (2021). Nuclei were diluted 5-fold in nuclei isolation buffer and stained with propidium iodide. 55,000 nuclei were sorted per sample using the S3e Cell Sorter (Bio-Rad, Hercules, USA) with FSC = 365 and SSC = 299. Nuclei were subsequently pelleted and Tagmentation and library amplification was done using the Zymo-seq ATAC library kit (D5458, Zymo Research, Irvine, USA). Sequencing was done with the Illumina Novaseq 6000 (150 bp paired-end, 20 million reads) at Baseclear (Leiden, the Netherlands).

### Bioinformatic analysis

ChIP-seq, RNA-seq, and ATAC-seq analyses were done using the nf-core ChIP-seq, ATAC-seq, or RNA-seq pipelines using Nextflow (Ewels et al., 2020), and reads were mapped to the TAIR10 version of the Arabidopsis genome. The threshold for peak identification by MACS2 in the nextflow pipeline was lowered to p <0.05 for ATAC-seq samples due to a low library complexity in these samples. For ChIP, reads were annotated in R using the ChIPSeeker package (Wang et al., 2022; Yu et al., 2015), and differential expression analysis for RNA-seq and differential accessibility analysis for ATAC-seq was done using the DESeq2 package (Love et al., 2014). Differentially expressed genes were calculated by comparing the DEX-treated *p35S:AHL15-GR* condition with the three control conditions (*p35S:AHL15-GR* + mock, Col-0 + mock and Col-0 + DEX), as the difference between control conditions was sufficiently small to exclude a strong effect of DEX on gene expression in Col-0 (Additional file 1: Figure S7). For ATAC-seq, mock- and DEX-treated samples at the same time point were compared. Reads were associated with genomic features using the Araport11 version of the Arabidopsis genome annotation.

Heatmaps of ChIP-seq reads were generated using deepTools (Ramírez et al., 2016). To annotate ChIP-seq peaks over genomic features, we used BEDTools (Quinlan and Hall, 2010) to categorize the TAIR10 genome into non-overlapping regions of the following categories: promoters (0-3 kb upstream of TSS), TES (0-1 kb downstream of TES), TES and promoter (regions where promoters and TES of adjacent genes overlapped), exon, intron, 5’ UTR, 3’ UTR, and intergenic (regions that are not covered by any of the other categories). Overlap between ChIP-seq and RNA-seq datasets was calculated in R, and ChIP-seq peaks shared between all AHL15 datasets, shared between 4 of the 5 AHL15 datasets, or unique to the overexpression datasets were identified using R and exported as BED files for further comparisons. Additionally, the ChIP peak coordinates of one *AHL15* overexpression ChIP-seq dataset were shuffled over the Arabidopsis genome to generate a set of random peak coordinates as a control. Motif enrichment analyses and de novo motif finding were done using the findMotifsGenome.pl function in HOMER2 with masking and a region size set to 50 bp (Heinz et al., 2010).

For the analysis of ChIP-seq data of epigenetic marks, raw data of epigenetic markers in wild type plants generated by Bourguet et al. (2021) was downloaded from GEO and reanalysed with the nf-core ChIP-seq pipeline. For generating profile plots of the various epigenetic marks over Arabidopsis genes or the AHL15 peaks, bigwig files of individual replicates were first normalized to input using a log2 normalization method and subsequently averaged before calculation of the count matrices and profile plotting with deepTools.

Gene identifiers of self-looping genes identified by Liu et al. (2016) (extracted from the supplementary data files) were compared with the annotated AHL15 ChIP-seq and RNA-seq datasets to identify which self-looping genes were bound by AHL15 and/or differentially expressed in AHL15-GR plants after DEX treatment. The proportion of self-looping genes in each category (unbound, bound & upregulated, bound & downregulated, bound but not differentially expressed) was compared between native and overexpression conditions using the χ^2^ test.

## Additional files

**Additional file 1: Figure S1:** Phenotype of 31 day-old *p35S:3xFLAG-AHL15* plants right before harvesting for ChIP. **Figure S2:** HOMER *de novo* motif finding results for overexpression-only peaks. **Figure S3:** HOMER de novo motif finding results for semi-shared peaks. **Figure S4:** HOMER de novo motif finding results calculated based on all AHL15 peaks (shared, semi-shared, and overexpression-only). **Figure S5:** HOMER *de novo* motif finding results using the 1000 highest-scoring shared AHL15 ChIP-seq peaks. **Figure S6:** HOMER *de novo* motif finding results using the AHL15-bound upregulated or AHL15-bound downregulated genes. **Figure S7:** Principal component analysis of RNA-seq datasets. **Figure S8:** Enrichment of ATAC-seq peaks over 3xFLAG-AHL15 ChIP-seq peaks showing that AHL15 binds DNA regions with reduced chromatin accessibility. **Figure S9:** Enrichment of H1, H2A, H2Z.X, H2A.Z, H3K9me1, H3K9me2, and H3K927me1 over 3xFLAG-AHL15 ChIP-seq peaks shows that AHL15 binding sites are characterized by a depletion of epigenetic marks. **Table S1:** HOMER *de novo* motif finding results using AHL15-bound self-looping genes.

**Additional file 2:** ChIP-seq peaks of 3xFLAG-AHL15 in native and overexpression conditions.

**Additional file 3:** HOMER2-identified motifs.

**Additional file 4:** Differential expression data of RNA-seq in *p35S:AHL15-GR* seedlings upon 8h DEX treatment.

**Additional file 5:** Genes that are bound by AHL15 and differentially expressed in *p35S:AHL15-GR* seedlings upon DEX treatment.

**Additional file 6:** ATAC-seq peaks in *p35S:AHL15-GR* seedlings after 30min, 1h, or 8h DEX or Mock treatment.

**Additional file 7:** Self-looping genes that are bound by AHL15.

## Declarations

### Ethics approval and consent to participate

Not applicable

### Consent for publication

Not applicable

### Availability of data and materials

All raw sequencing data has been deposited on SRA (Bioproject PRJNA1301884). The results of the analyses supporting the conclusions of this article are included within the article and its additional files as .xlsx files. Scripts used for the analyses and plant materials used for this study are available from Remko Offringa on reasonable request.

### Author contributions

TL and RO conceived and designed the study, with input from JC. TL performed all experiments. Data analysis was done by TL with help from JC. The manuscript was written by TL with input from JC and RO. All authors read and approved the manuscript.

## Acknowledgements

We thank Jan Vink, Altay Temel, Ward de Winter and Mariel Lavrijsen for technical support and Alex Bos, Marjolein Crooijmans and Dennis Claessen for help with nuclear sorting for ATAC-seq. We thank Salma Balazadeh for sharing her ChIP-seq protocol and Maike Stam for her advice on the ATAC-seq experiment.

## Literature

Aravind L. 1998. AT-hook motifs identified in a wide variety of DNA-binding proteins. Nucleic Acids Res. 26:4413–4421. DOI: 10.1093/nar/26.19.4413

Bourguet P, Picard CL, Yelagandula R, Pélissier T, Lorković ZJ, Feng S, Pouch-Pélissier MN, Schmücker A, Jacobsen SE, Berger F, Mathieu O. 2021. The histone variant H2A.W and linker histone H1 co-regulate heterochromatin accessibility and DNA methylation. Nat. Commun. 12:1–12. DOI: 10.1038/s41467-021-22993-5, PMID: 33976212

Charbonnel C, Rymarenko O, Ines O Da, Benyahya F, White CI, Butter F, Amiard S. 2018. The linker histone GH1-HMGA1 is involved in telomere stability and DNA damage repair. Plant Physiol. 177:311–327. DOI: 10.1104/PP.17.01789, PMID: 29622687

Chouaref J, de Boer E, Fransz P, Stam M. 2018. Protocol for Chromatin Immunoprecipitation of Meiotic-Stage-Specific Tomato Anthers. Curr. Protoc. Plant Biol. 3:e20074. DOI: 10.1002/cppb.20074

Clough SJ, Bent AF. 1998. Floral dip: a simplified method for Agrobacterium-mediated transformation of Arabidopsis thaliana. Plant J. 16:735–743. DOI: 10.1046/J.1365-313X.1998.00343.X, PMID: 10069079

den Dulk-Ras A, Hooykaas PJJ. 1995. Electroporation of Agrobacterium tumefaciens. Plant Cell Electroporation And Electrofusion Protocols. Humana Press. p. 63–72. DOI: 10.1385/0-89603-328-7:63

Ewels PA, Peltzer A, Fillinger S, Patel H, Alneberg J, Wilm A, Garcia MU, Di Tommaso P, Nahnsen S. 2020. The nf-core framework for community-curated bioinformatics pipelines. Nat. Biotech. 38:276–278. DOI: 10.1038/s41587-020-0439-x, PMID: 32055031

Favero DS, Kawamura A, Shibata M, Takebayashi A, Jung JH, Suzuki T, Jaeger KE, Ishida T, Iwase A, Wigge PA, Neff MM, Sugimoto K. 2020. AT-Hook transcription factors restrict petiole growth by antagonizing PIFs. Curr. Biol. 30:1454–1466. DOI: 10.1016/j.cub.2020.02.017, PMID: 32197081

Fletcher JC, Brand U, Running MP, Simon R, Meyerowitz EM. 1999. Signaling of cell fate decisions by CLAVATA3 in Arabidopsis shoot meristems. Science 283:1911–1914. DOI: 10.1126/SCIENCE.283.5409.1911/SUPPL_FILE/987103S3_THUMB.GIF, PMID: 10082464

Franco-Zorrilla JM, López-Vidriero I, Carrasco JL, Godoy M, Vera P, Solano R. 2014. DNA-binding specificities of plant transcription factors and their potential to define target genes. Proceedings of the National Academy of Sciences of the United States of America 111:2367–2372. DOI: 10.1073/pnas.1316278111, PMID: 24477691

Fujimoto S, Matsunaga S, Yonemura M, Uchiyama S, Azuma T, Fukui K. 2004. Identification of a novel plant MAR DNA binding protein localized on chromosomal surfaces. Plant Molecular Biology 56:225–239. DOI: 10.1007/s11103-004-3249-5

García-Heras F, Padmanabhan S, Murillo FJ, Elías-Arnanz M. 2009. Functional equivalence of HMGA- and histone H1-like domains in a bacterial transcriptional factor. PNAS 106:13546–13551. DOI: 10.1073/pnas.0902233106, PMID: 19666574

Heinz S, Benner C, Spann N, Bertolino E, Lin YC, Laslo P, Cheng JX, Murre C, Singh H, Glass CK. 2010. Simple combinations of lineage-determining transcription factors prime cis-regulatory elements required for macrophage and B cell identities. Molecular cell 38:576–89. DOI: 10.1016/j.molcel.2010.05.004, PMID: 20513432

Ji J, Strable J, Shimizu R, Koenig D, Sinha N, Scanlon MJ. 2010. WOX4 promotes procambial development. Plant Physiol. 152:1346–1356. DOI: 10.1104/PP.109.149641

Karami O, Henkel C, Offringa R. 2022. The effect of short-term activation of AHL15 on long-term plant developmental change and transcriptome profile. bioRxiv 2022.09.05.506087. DOI: 10.1101/2022.09.05.506087

Karami O, Rahimi A, Khan M, Bemer M, Hazarika RR, Mak P, Compier M, van Noort V, Offringa R. 2020. A suppressor of axillary meristem maturation promotes longevity in flowering plants. Nat. Plants 6:368–376. DOI: 10.1038/s41477-020-0637-z, PMID: 32284551

Karami O, Rahimi A, Mak P, Horstman A, Boutilier K, Compier M, van der Zaal B, Offringa R. 2021. An Arabidopsis AT-hook motif nuclear protein mediates somatic embryogenesis and coinciding genome duplication. Nature Communications 12:2508. DOI: 10.1038/s41467-021-22815-8

Karimi M, Depicker A, Hilson P. 2007. Recombinational Cloning with Plant Gateway Vectors. Plant Physiol. 145:1144–1154. DOI: 10.1104/PP.107.106989, PMID: 18056864

Kaufmann K, Muiño JM, Østerås M, Farinelli L, Krajewski P, Angenent GC. 2010. Chromatin immunoprecipitation (ChIP) of plant transcription factors followed by sequencing (ChIP-SEQ) or hybridization to whole genome arrays (ChIP-CHIP). Nat. Protoc. 5:457–472. DOI: 10.1038/nprot.2009.244, PMID: 20203663

Kim HB, Oh CJ, Park YC, Lee Y, Choe S, An CS, Choi SB. 2011. Comprehensive analysis of AHL homologous genes encoding AT-hook motif nuclear localized protein in rice. BMB Reports 44:680–685. DOI: 10.5483/BMBRep.2011.44.10.680

Klosterman SJ, Hadwiger LA. 2002. Plant HMG proteins bearing the AT-hook motif. Plant Science 162:855–866. DOI: 10.1016/S0168-9452(02)00056-0

Kumar A, Singh S, Mishra A. 2023. Genome-wide identification and analyses of the AHL gene family in rice (Oryza sativa). 3 Biotech 13. DOI: 10.1007/s13205-023-03666-0

Lee K, Seo PJ. 2017. Coordination of matrix attachment and ATP-dependent chromatin remodeling regulate auxin biosynthesis and Arabidopsis hypocotyl elongation. PLoS ONE 12:1–19. DOI: 10.1371/journal.pone.0181804

Lim PO, Kim Y, Breeze E, Koo JC, Woo HR, Ryu JS, Park DH, Beynon J, Tabrett A, Buchanan-Wollaston V, Nam HG. 2007. Overexpression of a chromatin architecture-controlling AT-hook protein extends leaf longevity and increases the post-harvest storage life of plants. The Plant Journal 52:1140–1153. DOI: 10.1111/j.1365-313X.2007.03317.x

Lin L, Nakano H, Nakamura S, Uchiyama S, Fujimoto S, Matsunaga S, Kobayashi Y, Ohkubo T, Fukui K. 2007. Crystal structure of Pyrococcus horikoshii PPC protein at 1.60 Å resolution. Proteins: Structure, Function, and Bioinformatics 67:505–507. DOI: 10.1002/PROT.21270, PMID: 17295322

Liu C, Wang C, Wang G, Becker C, Zaidem M, Weigel D. 2016. Genome-wide analysis of chromatin packing in Arabidopsis thaliana at single-gene resolution. Genome Res. 26:1057–1068. DOI: 10.1101/GR.204032.116/-/DC1, PMID: 27225844

Liu J, Chen Z, Sun L, Huang T, Chen Y, Li X, Liu H, Huang X, Peng Y, Feng B. 2026. An AT-hook motif nuclear protein AHL13 interacts with Poly(ADP-ribose) to regulate Arabidopsis immunity. Plant Science 365:113021. DOI: 10.1016/J.PLANTSCI.2026.113021

Love MI, Huber W, Anders S. 2014. Moderated estimation of fold change and dispersion for RNA-seq data with DESeq2. Genome Biology 15:550. DOI: 10.1186/s13059-014-0550-8

Luden T, Amakorová P, Novák O, Balazadeh S, Offringa R. 2025. The plant longevity gene AHL15 delays leaf senescence by repressing ORESARA1 and cytokinin degradation. bioRxiv 2025.11.23.689984. DOI: 10.1101/2025.11.23.689984

Machaj G, Grzebelus D. 2021. Characteristics of the at-hook motif containing nuclear localized (Ahl) genes in carrot provides insight into their role in plant growth and storage root development. Genes 12:764. DOI: 10.3390/genes12050764, PMID: 34069875

Murashige T, Skoog F. 1962. A Revised Medium for Rapid Growth and Bio Assays with Tobacco Tissue Cultures. Physiologia Plantarum 15:473–497. DOI: 10.1111/J.1399-3054.1962.TB08052.X

Ng KH, Yu H, Ito T. 2009. AGAMOUS controls GIANT KILLER, a multifunctional chromatin modifier in reproductive organ patterning and differentiation. PLoS Biol. 7:e1000251. DOI: 10.1371/journal.pbio.1000251

Ozturk N, Singh I, Mehta A, Braun T, Barreto G. 2014. HMGA proteins as modulators of chromatin structure during transcriptional activation. Front. Cell Dev. Biol. 2:1–9. DOI: 10.3389/fcell.2014.00005

Quinlan AR, Hall IM. 2010. BEDTools: a flexible suite of utilities for comparing genomic features. Bioinformatics 26:841. DOI: 10.1093/BIOINFORMATICS/BTQ033, PMID: 20110278

Rahimi A, Karami O, Balazadeh S, Offringa R. 2022a. miR156 -independent repression of the ageing pathway by longevity-promoting AHL proteins in Arabidopsis. New Phytol. 235:2424–2438. DOI: 10.1111/NPH.18292

Rahimi A, Karami O, Lestari AD, de Werk T, Amakorová P, Shi D, Novák O, Greb T, Offringa R. 2022b. Control of cambium initiation and activity in Arabidopsis by the transcriptional regulator AHL15. Curr. Biol. 32:1764–1775. DOI: 10.1016/J.CUB.2022.02.060, PMID: 35294866

Ramírez F, Ryan DP, Grüning B, Bhardwaj V, Kilpert F, Richter AS, Heyne S, Dündar F, Manke T. 2016. deepTools2: a next generation web server for deep-sequencing data analysis. Nucleic Acids Research 44:W160–W165. DOI: 10.1093/nar/gkw257

Street IH, Shah PK, Smith AM, Avery N, Neff MM. 2008. The AT-hook-containing proteins SOB3/AHL29 and ESC/AHL27 are negative modulators of hypocotyl growth in Arabidopsis. Plant Journal 54:1–14. DOI: 10.1111/j.1365-313X.2007.03393.x

Tayengwa R, Sharma Koirala P, Pierce CF, Werner BE, Neff MM. 2020. Overexpression of AtAHL20 causes delayed flowering in Arabidopsis via repression of FT expression. BMC Plant Biol. 20:559. DOI: 10.1186/s12870-020-02733-5, PMID: 33308168

Wang FX, Shang GD, Wu LY, Mai YX, Gao J, Xu ZG, Wang JW. 2021. Protocol for assaying chromatin accessibility using ATAC-seq in plants. STAR Protocols 2. DOI: 10.1016/j.xpro.2020.100289, PMID: 33532736

Wang Q, Li M, Wu T, Zhan L, Li L, Chen M, Xie W, Xie Z, Hu E, Xu S, Yu G. 2022. Exploring Epigenomic Datasets by ChIPseeker. Curr. protoc. 2:e585. DOI: 10.1002/CPZ1.585, PMID: 36286622

Xiao C, Chen F, Yu X, Lin C, Fu YF. 2009. Over-expression of an AT-hook gene, AHL22, delays flowering and inhibits the elongation of the hypocotyl in Arabidopsis thaliana. Plant Mol. Biol. 71:39–50. DOI: 10.1007/S11103-009-9507-9/FIGURES/6, PMID: 19517252

Xiong SX, Zeng QY, Hou JQ, Hou LL, Zhu J, Yang M, Yang ZN, Lou Y. 2020. The temporal regulation of TEK contributes to pollen wall exine patterning. PLoS Genet. 16:e1008807. DOI: 10.1371/journal.pgen.1008807, PMID: 32407354

Xu L, Zheng S, Witzel K, Van De Slijke E, Baekelandt A, Mylle E, Van Damme D, Cheng J, De Jaeger G, Inzé D, Jiang H. 2024. Chromatin attachment to the nuclear matrix represses hypocotyl elongation in Arabidopsis thaliana. Nat. Commun. 15:1286. DOI: 10.1038/s41467-024-45577-5

Xu Y, Wang Y, Stroud H, Gu X, Sun B, Gan ES, Ng KH, Jacobsen SE, He Y, Ito T. 2013. A matrix protein silences transposons and repeats through interaction with retinoblastoma-associated proteins. Curr. Biol. 23:345–350. DOI: 10.1016/j.cub.2013.01.030

Yu G, Wang LG, He QY. 2015. ChIPseeker: an R/Bioconductor package for ChIP peak annotation, comparison and visualization. Bioinformatics 31:2382–2383. DOI: 10.1093/BIOINFORMATICS/BTV145, PMID: 25765347

Yun J, Kim Y-S, Jung J-H, Seo PJ, Park C-M. 2012. The AT-hook motif-containing protein AHL22 regulates flowering initiation by modifying FLOWERING LOCUS T chromatin in Arabidopsis. J. Biol. Chem. 287:15307–15316. DOI: 10.1074/jbc.M111.318477

Zhao B, Xi Y, Kim J, Sung S. 2021. Chromatin architectural proteins regulate flowering time by precluding gene looping. Sci. Adv. 7:3097–3108. DOI: 10.1126/SCIADV.ABG3097/SUPPL_FILE/SCIADV.ABG3097_SM.PDF, PMID: 34117065

Zhao J, Favero DS, Peng H, Neff MM. 2013. Arabidopsis thaliana AHL family modulates hypocotyl growth redundantly by interacting with each other via the PPC/DUF296 domain. PNAS 110:E4688–E4697. DOI: 10.1073/pnas.1219277110

Zhao J, Favero DS, Qiu J, Roalson EH, Neff MM. 2014. Insights into the evolution and diversification of the AT-hook Motif Nuclear Localized gene family in land plants. BMC Plant Biol. 14:1–19. DOI: 10.1186/s12870-014-0266-7

Zhou J, Wang X, Lee Jung Youn, Lee Ji Young. 2013. Cell-to-Cell Movement of Two Interacting AT-Hook Factors in Arabidopsis Root Vascular Tissue Patterning. Plant Cell 25:187–201. DOI: 10.1105/TPC.112.102210, PMID: 23335615

