## Additional files 1-7 for "AT-HOOK-MOTIF NUCLEAR LOCALIZED 15 extends plant longevity by binding at poorly accessible, epigenetic mark-depleted chromatin surrounding transcribed regions": Additional_file_1_supplemental_figures_revised.pdf

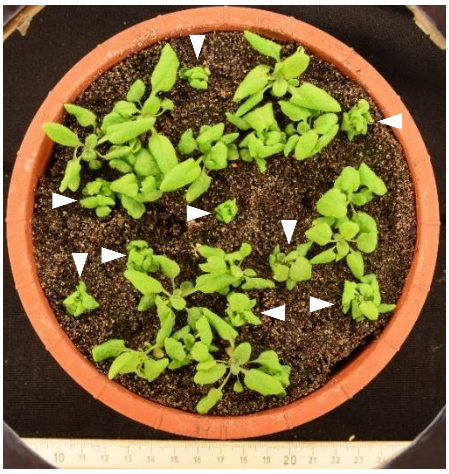

**Figure S1:** Phenotype of 31 day-old *p35S:3xFLAG-AHL15* plants right before harvesting for ChIP. White arrowheads mark plants with strong AHL15-induced delay of development that were used for *3xFLAG-AHL15* overexpression ChIP. Plants were grown in long day condition (16h light / 8h dark), and the scale on the bottom of the image is in cm.

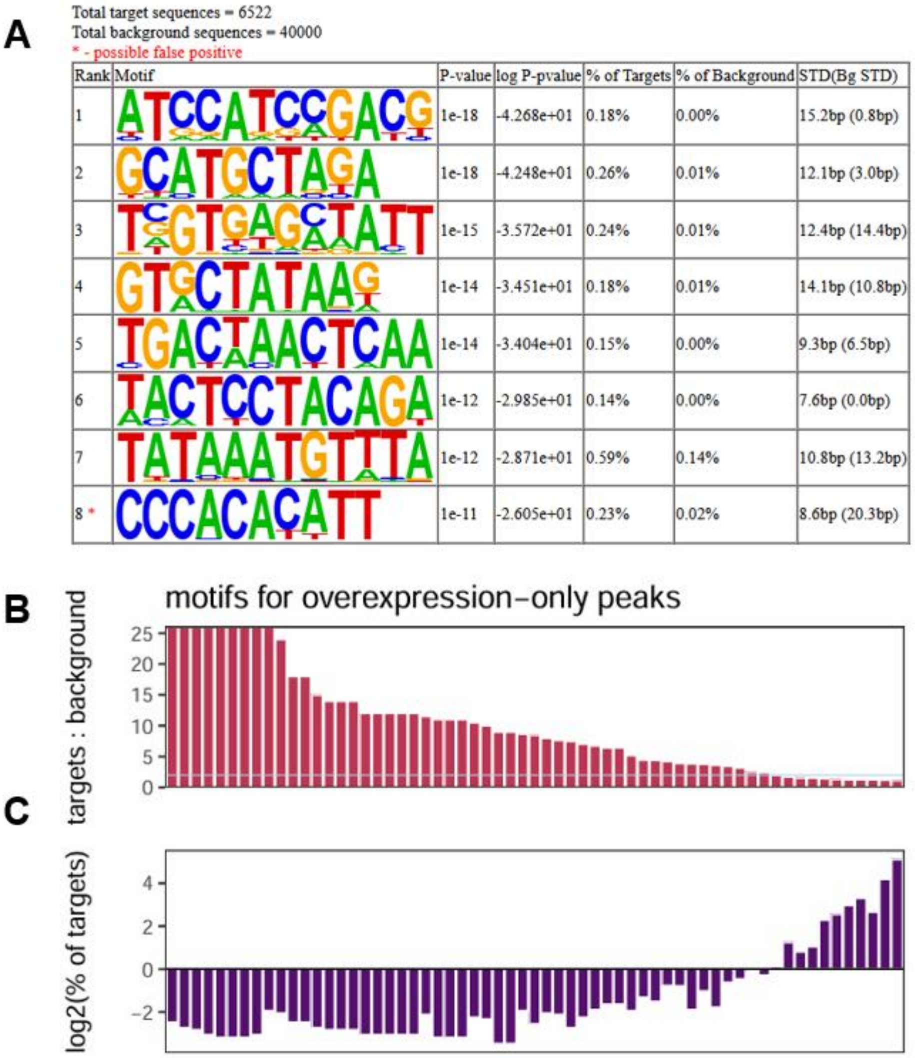

Total target sequences = 6522
Total background sequences = 40000
* - possible false positive

| Rank | Motif | P-value | log P-pvalue | % of Targets | % of Background | STD(Bg STD) |
|---|---|---|---|---|---|---|
| 1 | ATCCATCCGACG | 1e-18 | -4.268e+01 | 0.18% | 0.00% | 15.2bp (0.8bp) |
| 2 | GCATGCTAGA | 1e-18 | -4.248e+01 | 0.26% | 0.01% | 12.1bp (3.0bp) |
| 3 | TCGTGAGCTATT | 1e-15 | -3.572e+01 | 0.24% | 0.01% | 12.4bp (14.4bp) |
| 4 | GTGCTATAAG | 1e-14 | -3.451e+01 | 0.18% | 0.01% | 14.1bp (10.8bp) |
| 5 | TGACTAACTCAA | 1e-14 | -3.404e+01 | 0.15% | 0.00% | 9.3bp (6.5bp) |
| 6 | TACTCCTACAGA | 1e-12 | -2.985e+01 | 0.14% | 0.00% | 7.6bp (0.0bp) |
| 7 | TATAAATGTTTA | 1e-12 | -2.871e+01 | 0.59% | 0.14% | 10.8bp (13.2bp) |
| 8 * | CCCACACATT | 1e-11 | -2.605e+01 | 0.23% | 0.02% | 8.6bp (20.3bp) |



**Figure S2**: HOMER de novo motif finding results for overexpression-only peaks. **A:** Table of the top 8 enriched motifs identified by HOMER2 within the set of overexpression-only peaks, with the p-value of motif enrichment, $\log_{10}$ of the p-value of motif enrichment, the percent of AHL15 peaks (targets) containing the motif, and the percent of background sequences containing the motif. **B:** The number of times each motif (61 in total) is identified by HOMER2 in overexpression-only AHL15 peaks divided by its number over the entire genome (expressed as %: 100% means the motif is almost only present in AHL15 peaks). **C:** The frequency with which each motif identified by HOMER2 occurs in overexpression-only AHL15 peaks, expressed as the Log2 value of the % of shared AHL15 targets containing the specific motif (occurrence of more than 1% gives a positive value, 1% gives a value of 0, and less than 1% results in a negative value). Motifs in figures **B** and **C** are arranged by descending target : background value.

**A**

Total target sequences = 4000
Total background sequences = 51667
* - possible false positive

| Rank | Motif | P-value | log P-pvalue | % of Targets | % of Background | STD(Bg STD) |
|---|---|---|---|---|---|---|
| 1 | AACATCAATCTT | 1e-19 | -4.398e+01 | 0.38% | 0.01% | 10.8bp (9.3bp) |
| 2 | TGTAAAAATTAA | 1e-15 | -3.683e+01 | 0.28% | 0.01% | 10.2bp (5.9bp) |
| 3 | AAATACAATTGC | 1e-14 | -3.269e+01 | 0.25% | 0.01% | 9.5bp (10.4bp) |
| 4 | TGCAGATGCCCT | 1e-14 | -3.263e+01 | 0.35% | 0.02% | 13.7bp (12.0bp) |
| 5 | AATATTTAAATT | 1e-13 | -3.124e+01 | 8.03% | 5.18% | 11.4bp (13.0bp) |
| 6 | TGTACCGCAT | 1e-13 | -3.017e+01 | 0.20% | 0.00% | 8.2bp (0.0bp) |
| 7 | GATCTAAGCAAA | 1e-13 | -3.017e+01 | 0.20% | 0.00% | 10.6bp (1.7bp) |
| 8 | TTAACTAGACAT | 1e-12 | -2.937e+01 | 0.28% | 0.01% | 12.7bp (14.0bp) |
| 9 | ATAAGGGTCATA | 1e-12 | -2.872e+01 | 0.25% | 0.01% | 9.2bp (10.2bp) |
| 10 | CGGTGAAATCCT | 1e-12 | -2.864e+01 | 0.23% | 0.01% | 11.5bp (7.6bp) |
| 11 * | CCTTAAACACCA | 1e-11 | -2.592e+01 | 0.25% | 0.01% | 10.4bp (5.8bp) |

**B**

**C**

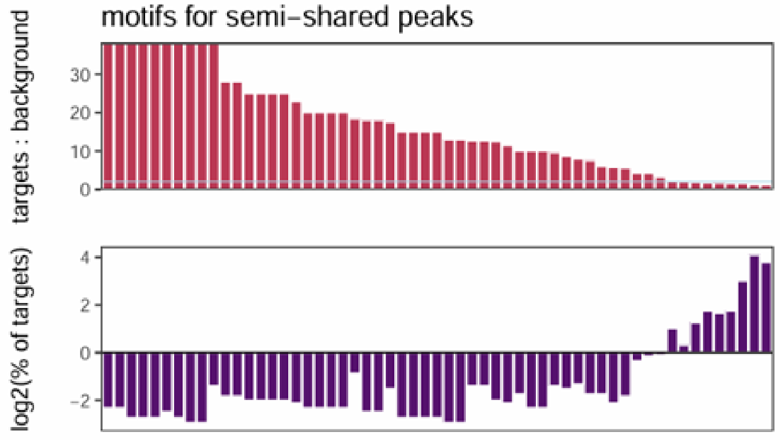


**Figure S3**: HOMER de novo motif finding results for semi-shared peaks. **A:** Table of the top 11 enriched motifs identified by HOMER2 within the set of semi-shared peaks, with the p-value of motif enrichment, $log_{10}$ of the p-value of motif enrichment, the percent of AHL15 peaks (targets) containing the motif, and the percent of background sequences containing the motif. **B:** The number of times each motif (57 in total) is identified by HOMER2 in semi-shared AHL15 peaks divided by its number over the entire genome (expressed as %: 100% means the motif is almost only present in AHL15 peaks). **C:** The frequency with which each motif identified by HOMER2 occurs in semi-shared AHL15 peaks, expressed as the Log2 value of the % of shared AHL15 targets containing the specific motif (occurrence of more than 1% gives a positive value, 1% gives a value of 0, and less than 1% results in a negative value). Motifs in figures **B** and **C** are arranged by descending target : background value.

**A**

Total target sequences = 12083
Total background sequences = 40333
* - possible false positive

| Rank | Motif | P-value | log P-pvalue | % of Targets | % of Background | STD(Bg STD) |
|---|---|---|---|---|---|---|
| 1 | AACATCAATCTT | 1e-23 | -5.519e+01 | 0.17% | 0.01% | 11.2bp (9.9bp) |
| 2 | TAGTTTACAC | 1e-18 | -4.155e+01 | 0.14% | 0.01% | 9.2bp (6.3bp) |
| 3 | TTAAACGGCTTT | 1e-18 | -4.155e+01 | 0.14% | 0.01% | 11.5bp (4.2bp) |
| 4 | TCAAAGTACATC | 1e-17 | -4.087e+01 | 0.16% | 0.01% | 13.7bp (10.2bp) |
| 5 | GAAATATATCCC | 1e-17 | -3.948e+01 | 0.18% | 0.02% | 11.2bp (7.8bp) |
| 6 | TGCAATCTCG | 1e-16 | -3.828e+01 | 0.13% | 0.01% | 9.1bp (19.7bp) |
| 7 | TGGAGATGCTCT | 1e-16 | -3.789e+01 | 0.15% | 0.01% | 12.5bp (15.6bp) |
| 8 | AATGGTACAA | 1e-15 | -3.573e+01 | 0.24% | 0.03% | 14.2bp (13.1bp) |
| 9 | TAAAAGAATACG | 1e-15 | -3.506e+01 | 0.12% | 0.01% | 13.0bp (8.1bp) |
| 10 | GAACTAGATTTC | 1e-15 | -3.506e+01 | 0.12% | 0.01% | 8.1bp (1.6bp) |
| 11 | AGTTATACTAAG | 1e-15 | -3.496e+01 | 0.14% | 0.01% | 11.2bp (10.9bp) |
| 12 | AGGTAATATA | 1e-14 | -3.322e+01 | 0.41% | 0.11% | 12.1bp (15.7bp) |
| 13 | AAACTTTTTCAG | 1e-14 | -3.269e+01 | 0.18% | 0.02% | 10.5bp (11.0bp) |
| 14 | TCACGATAATTT | 1e-13 | -3.192e+01 | 0.12% | 0.01% | 12.3bp (17.4bp) |
| 15 | AATCATATTTAA | 1e-13 | -3.177e+01 | 0.16% | 0.02% | 10.5bp (8.9bp) |
| 16 | TGGACGGG | 1e-13 | -3.173e+01 | 0.21% | 0.03% | 12.8bp (14.7bp) |
| 17 | TGCCAGCC | 1e-13 | -3.170e+01 | 3.14% | 2.08% | 12.2bp (13.9bp) |
| 18 | GAATATACATTT | 1e-12 | -2.779e+01 | 0.13% | 0.01% | 13.0bp (3.4bp) |
| 19 | TAAATCCGTCAC | 1e-12 | -2.778e+01 | 0.18% | 0.03% | 12.6bp (10.0bp) |
| 20 * | AATGGCCA | 1e-11 | -2.663e+01 | 0.22% | 0.04% | 12.9bp (12.6bp) |

**B**

**C**

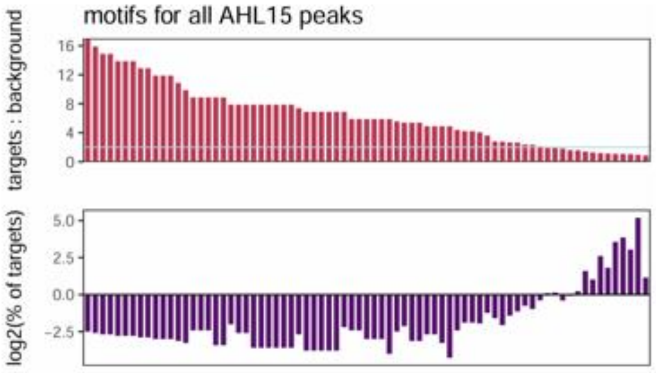


**Figure S4:** HOMER de novo motif finding results calculated based on all AHL15 peaks (shared, semi-shared, and overexpression-only). **A:** Table of the top 20 enriched motifs identified by HOMER2 within all AHL15 peaks, with the p-value of motif enrichment, $log_{10}$ of the p-value of motif enrichment, the percent of AHL15 peaks (targets) containing the motif, and the percent of background sequences containing the motif. **B:** The number of times each motif (75 in total) is identified by HOMER2 in all AHL15 peaks divided by its number over the entire genome (expressed as %: 100% means the motif is almost only present in AHL15 peaks). **C:** The frequency with which each motif identified by HOMER2 occurs in all AHL15 peaks, expressed as the Log2 value of the % of shared AHL15 targets containing the specific motif (occurrence of more than 1% gives a positive value, 1% gives a value of 0, and less than 1% results in a negative value). Motifs in figures **B** and **C** are arranged by descending target : background value.

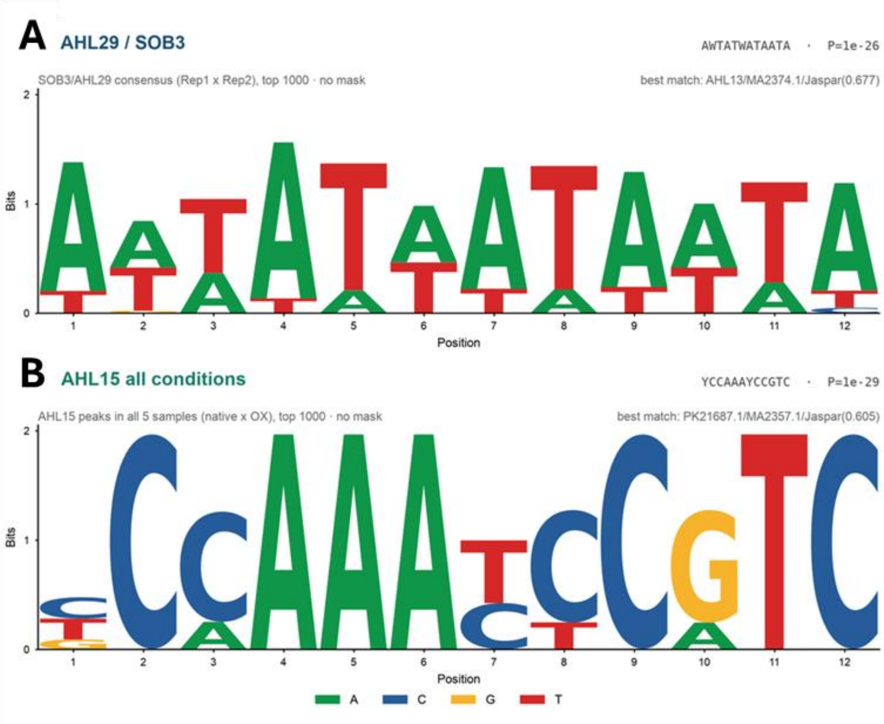


**Figure S5**: HOMER *de novo* motif finding results calculated based on **A** the 1000 highest-scoring ChIP-seq peaks of AHL29 (data taken from Favero et al., 2020, current biology) or **B** the 1000 highest-scoring shared AHL15 ChIP-seq peaks.

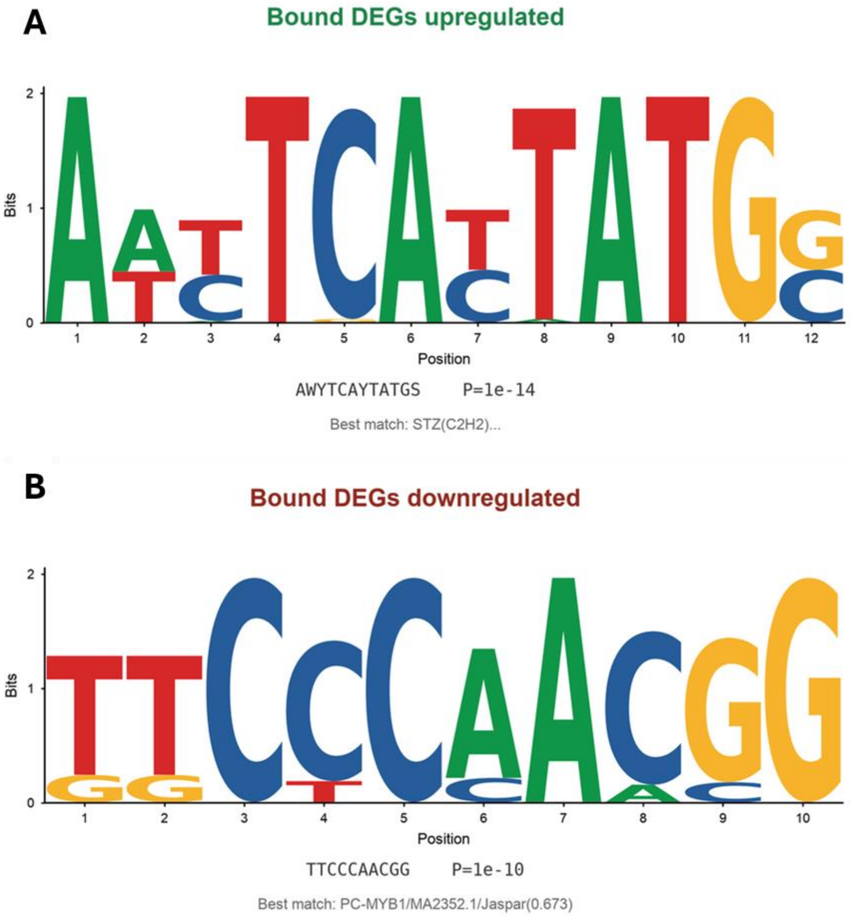


**Figure S6**: HOMER *de novo* motif finding results in AHL15-bound differentially expressed genes. **A:** AHL15-bound upregulated genes, **B:** AHL15-bound downregulated genes.

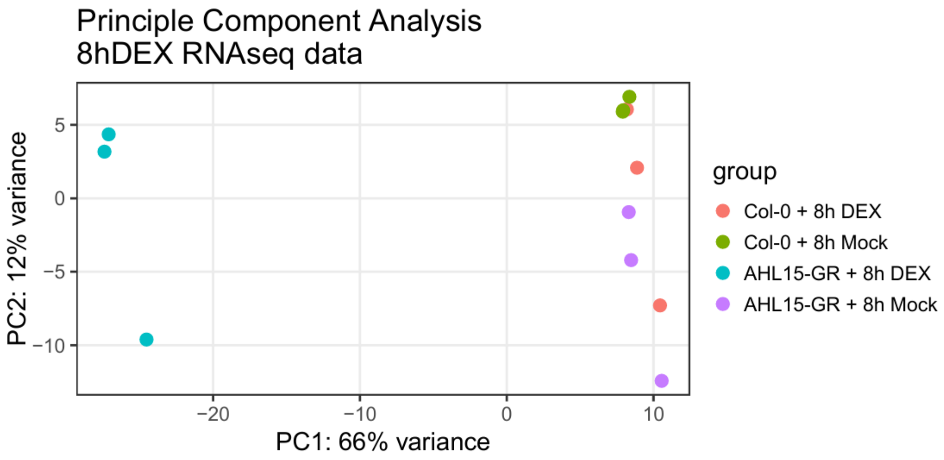


**Figure S7:** Principal component analysis of RNA-seq datasets. RNA-seq was performed on RNA isolated from 10 day-old wild-type (Col-0) or *p35S:AHL15-GR* seedlings that mock or DEX treated for 15 minutes. Per replicate (n=3) shoot tissue of five plants was harvested at 8 hours after treatment for RNA isolation and sequencing.

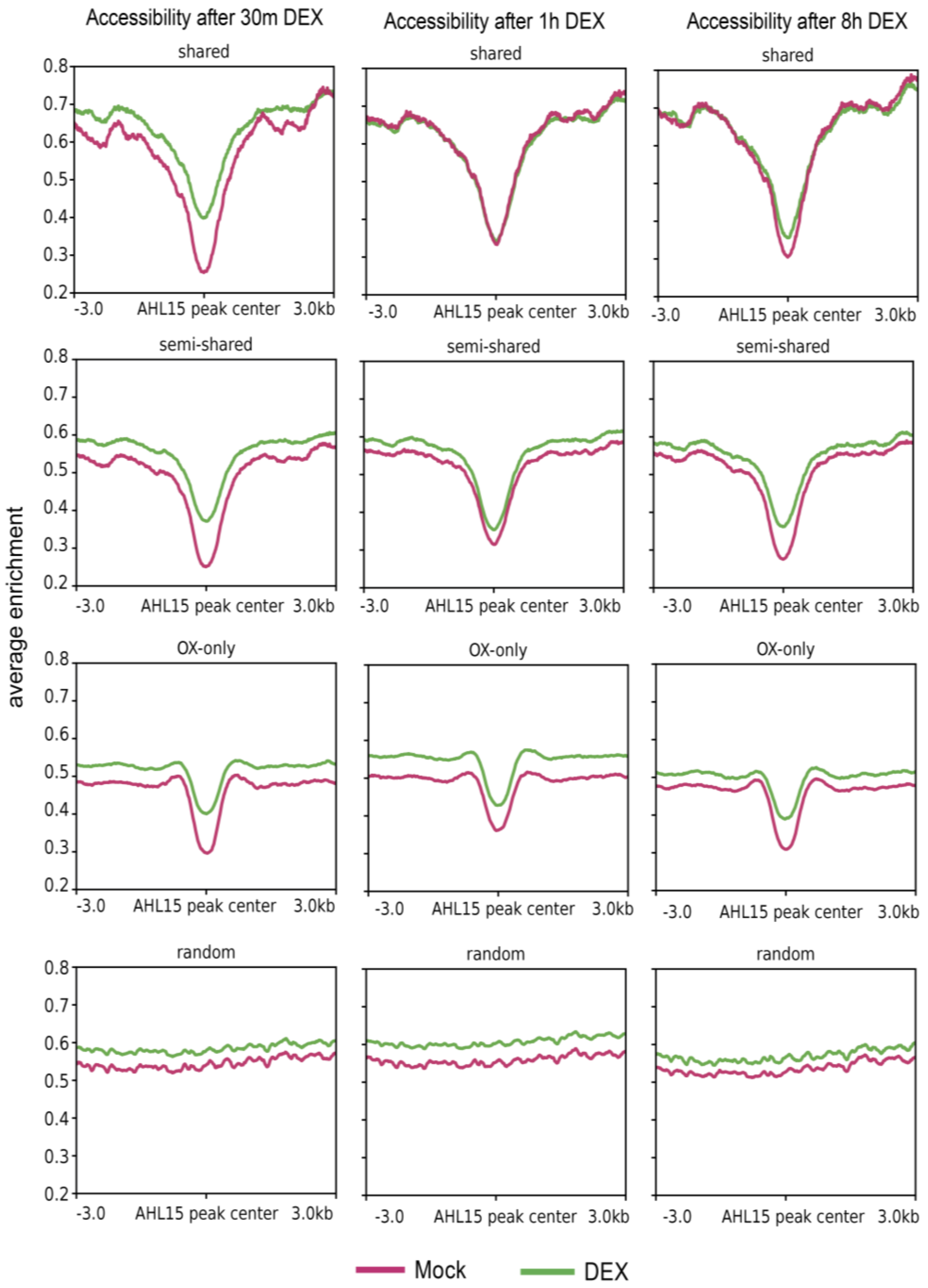


**Figure S8:** Enrichment of ATAC-seq peaks over 3xFLAG-AHL15 ChIP-seq peaks showing that AHL15 binds DNA regions with reduced chromatin accessibility. ATAC-seq reads from *p35S:AHL15-GR* plants treated with DEX or mock for 30 minutes, 1 hour or 8 hours plotted over three classes of AHL15 ChIP-seq peaks presented in Figure 1C: shared (identified in native and overexpression conditions, n = 1948), semi-shared (identified in one native sample and in all overexpression samples, n = 2906), and OX-only (identified in all three overexpression samples, n = 4999). Random genomic coordinates with similar size distribution as the ChIP-seq peaks were used as a negative control (random, n = 15221 regions).

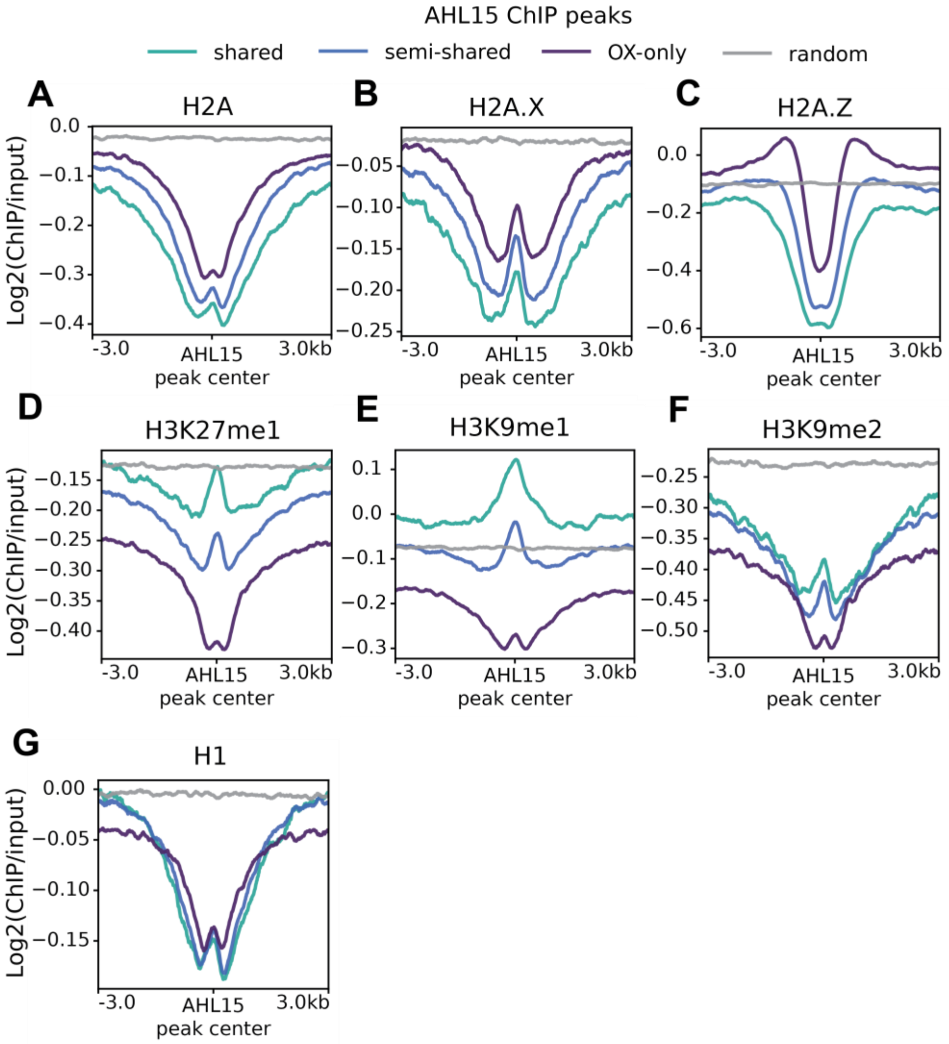


**Figure S9:** Enrichment of H1, H2A, H2Z.X, H2A.Z, H3K9me1, H3K9me2, and H3K927me1 over 3xFLAG-AHL15 ChIP-seq peaks shows that AHL15 binding sites are characterized by a depletion of epigenetic marks. **A-G:** ChIP-seq profiles for the epigenetic marks H2A (**A**), H2A.X (**B**), H2A.Z **(C**), H3K27me1 (**D**), H3K9me1 (**E**), H3K9me2 (**F**), and H1 (**G**) plotted over three classes of AHL15 ChIP-seq peaks presented in Figure 1C: shared (identified in native and overexpression conditions, n = 1948), semi-shared (identified in one native sample and in all overexpression samples, n = 2906), and OX-only (identified in all three overexpression samples, n = 4999). Random genomic coordinates with similar size distribution as the ChIP-seq peaks were used as a negative control (random, n = 15221 regions).

**Table S1:** HOMER *de novo* motif finding results using the AHL15-bound and differentially expressed self-looping genes with the closest known matching motifs.

| Set | Pooled peaks | Top motif | Enrichment P | Closest public match |
|---|---|---|---|---|
| Self-loop native up | 74 | AAAATTGAATCG | 1.00E-08 | AT1G20910(ARID)/col-AT1G20910-DAP-Seq |
| Self-loop native down | 50 | TATTATTTTA | 1.00E-06 | SPL15B/MA2432.1/Jaspar(0.704) |
| Self-loop OX up | 387 | AAMTAGWKAATG | 1.00E-18 | HDG1/MA1369.2/Jaspar(0.622) |
| Self-loop OX down | 345 | GATKCGACGMAG | 1.00E-17 | Replumless(BLH)/Arabidopsis-RPL.GFP-ChIP-Seq |
| Self-loop all-cond. up | 461 | TCATTAATCTGC | 1.00E-18 | ZHD10/MA1807.2/Jaspar(0.758) |
| Self-loop all-cond. down | 395 | CTTTTTATATTR | 1.00E-16 | POL012.1_TATA-Box/Jaspar(0.649) |
